# In-cell structures visualize human pre-ribosome assembly in the nucleolus

**DOI:** 10.64898/2026.06.01.729245

**Authors:** Xiaohan Zhao, Yuki Hayashi, Herman K.H. Fung, Sara Cuylen-Häring, Julia Mahamid, Christoph W. Müller

## Abstract

Ribosome biogenesis is an essential multi-step process. In eukaryotes, it is orchestrated within the nucleolus, a hallmark multi-layered compartment of the nucleus. Structures of pre-ribosomes have been characterized *in vitro*, but their assembly pathways within human nucleoli have not been described. Here, we used cryogenic correlative light and electron tomography to visualize the molecular landscapes within HeLa cell nucleoli and obtained in-cell structural snapshots of both ribosomal subunit precursors, the SSU processome and the pre-60S. These recapitulate major states resolved previously *in vitro* while additionally revealing critical interaction partners, including the RNA exosome, rixosome, and nuclear export receptor CRM1-RanGTP. We further show how pre-ribosome assembly is altered upon RNA polymerase I inhibition. Our study combines molecular structures with cellular context to elucidate the spatiotemporal assembly pathways and regulation of human ribosome biogenesis.

## Main Text

Ribosome biogenesis is one of the most active processes in cells, and, in eukaryotes, spans three subcellular compartments: the nucleolus, the nucleus, and the cytoplasm (*1*). Ribosomal RNA (rRNA) transcription and assembly of early ribosome precursors (pre-ribosomes) are orchestrated in the nucleolus (*2, 3*) a membrane-less, multi-layered compartments composed of fibrillar centers (FCs), dense fibrillar components (DFCs) and granular components (GCs) (*4*). Transcription of rRNA occurs at the interface of the FC and DFC (*5*). Here, RNA polymerase I (Pol I) initially synthesizes a large 47S pre-rRNA transcript comprising the 18S rRNA, the major scaffold of the small subunit (SSU), and 28S and 5.8S rRNA, two of three large subunit (LSU) rRNAs. These regions are joined by internal transcribed spacers (ITS1 and ITS2) and flanked by external transcribed spacers (5′ ETS and 3′ ETS). Nascent rRNA transcripts are initially modified and processed in the DFC, and subsequently further processed and assembled into pre-ribosomes within the GC (*5*–*7*). This process involves a large number of ribosomal proteins and transiently associated assembly factors (*8*). Studies by single-particle cryo-electron microscopy (cryo-EM) provided high-resolution structures of pre-ribosome assembly intermediates, offering insights into the mechanisms underlying the maturation of the human ribosomal subunits (*9*–*13*). However, *in vitro* approaches inevitably lose the native cellular context. A complementary in-cell cryo-electron tomography (cryo-ET) study of nucleoli in *Chlamydomonas reinhardtii* revealed gradients of ribosome maturation, but the moderate resolution (∼25Å) of the reported pre-ribosomes reconstructions (*14*) limited a detailed understanding of their assembly pathway.

To gain deeper insight into human nucleolar ribosome biogenesis, we employed cryo-ET guided by correlative light and electron microscopy (CLEM) to investigate the molecular landscape within nucleoli in intact HeLa cells. This approach allowed us to localize nucleoli in cryo-fixed cells and obtain in-cell cryo-ET maps of both ribosomal subunit precursors, the SSU processome and the pre-60S, at 8.2 Å and 6.9 Å resolution, respectively. These represent the highest resolution in-cell structures resolved in the nucleus to date. Classification recapitulated two *in vitro* reported states of the human SSU processome and eight states similar to previously identified human pre-60S intermediates *in vitro*. In addition, we captured multiple novel transient intermediates. Finally, by perturbing the system with Pol I inhibitor actinomycin D (ActD), we visualized how pre-ribosome assembly is reconfigured in response to canonical nucleolar stress. By integrating high-resolution structural data with cellular context, we reconstruct the spatiotemporal assembly pathway of human pre-ribosomes and uncover new layers of regulation within the human nucleolus.

### Cryo-ET reveals molecular landscape of ribosome biogenesis in the human nucleolus

The three main compartments of the human nucleolus (FC, DFC and GC) were historically defined based on differences in their apparent textures in heavy-metal-stained transmission electron microscopy (TEM) images. In these images, the DFC appeared as the most electron-dense nucleolar compartment, whereas the FC appeared lighter and the GC showed a characteristic granular texture presumably due to the presence of pre-ribosomal particles (*15*). In live cells, these three nucleolar components can be also represented by the specific localization of different proteins; for example, upstream binding factor (UBF) localizes in the FC, fibrillarin (FBL) localizes in the DFC, and nucleophosmin 1 (NPM1) localizes in the GC (*16*) (fig. S1A). To locate the multi-layered nucleolus in cryo-TEM, we thus generated HeLa cell lines with fluorescently tagged proteins and developed a cryogenic CLEM workflow to guide cryo-ET data acquisition (Methods). Specifically, we registered multi-channel cryo-Airyscan fluorescence images of cryo-focused ion beam (cryo-FIB) prepared lamellae with their cryo-TEM maps to identify the locations of different nucleolar compartments on the lamellae and in the subsequently reconstructed tomograms (Fig. 1, A and B).

**Fig. 1.**
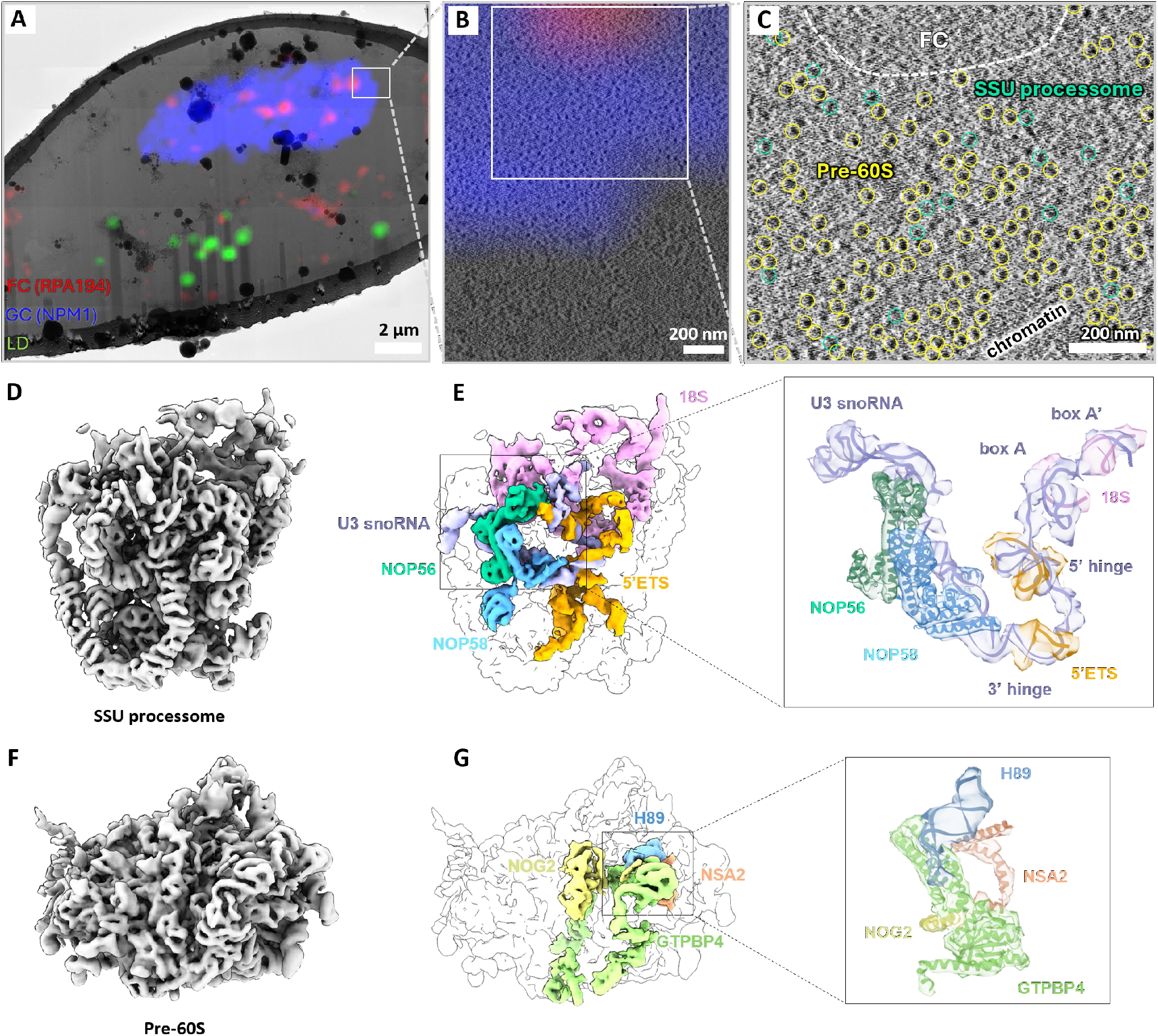
Pre-ribosome structures in human nucleoli. (**A** to **C**) Cryo-CLEM guided TEM data acquisition in a triple labeled HeLa cell line (mEGFP-NPM1, Halo-UBF (not shown here), and SNAP-RPA194 stained with JFX-SNAP646), corresponding tomograms with registered fluorescence, and denoised tomogram slice (*50*). SSU processome and pre-60S particles are indicated in green and yellow circles, respectively. LD: lipid droplet. Live stain by BODIPY 558/568, used for accurate registration of the CLEM data. (**D**) Cryo-ET consensus map of SSU processome, global resolution= 8.2 Å. (**E**) Segmented ribosomal RNA scaffold and protein factors in SSU processome. Zoom in shows local fitting of 18S rRNA, 5’ETS, U3 snoRNA, NOP56 and NOP58. Molecular models from PDB 7mq8. (**F**) Cryo-ET consensus map of pre-60S, global resolution= 6.9 Å. (**G**) Segmented 28S rRNA helix H89 and peripheral protein factors in pre-60S. Zoom in shows local fitting of H89, GTPB4, NSA2 and peptide of NOG2. Molecular models from PDB 8fla.

In contrast to the clear textural difference seen by room temperature TEM (*15*), we observed no obvious substructures of the nucleoli in either low magnification lamella maps (Fig. 1A) or high magnification reconstructed tomograms (Fig. 1B and fig. S1, B and C). Nevertheless, the particle distributions correlated with the fluorescence-defined nucleolar compartments. Significantly, large particles, presumably corresponding to SSU processome and pre-60S particles (Fig. 1C, circles), stood out from the background in the NPM1-fluorescent GC area. These two types of particles were picked and processed independently with subtomogram analysis (Methods), providing consensus maps for the SSU processome as the early biogenesis precursor of the small subunit (*17*) (Fig. 1, D and E) and for pre-60S representing the precursor of the large subunit (*9*) (Fig. 1, F and G). The local resolution of both consensus maps reached the Nyquist limit (6.85 Å) of the data (fig. S1D) and revealed secondary-structure level details (Fig. 1, E and G, and fig. S1E).

The densities provided structural information of the proteins and rRNAs, and the interactions between them, revealing key structural features of pre-ribosome assembly that are consistent with *in vitro* studies. For example, in the SSU processome, small nucleolar RNA U3 (U3 snoRNA) forms a central architectural anchor point to coordinate both the 5′ ETS via the 3′ and 5′ hinges and the 18S rRNA via the A and A′ boxes (*18*). The canonical Box C/D snoRNP members, NOP56 and NOP58, bind to the 3’ of U3 snoRNA via their Nop domains (*19, 20*) (Fig. 1E). In the pre-60S, the N-terminal domain of GTPB4 is inserted into the two strands of helix H89 of the 28S rRNA, placing H89 in a different position compared to its mature form. NSA2 and NOG2 bind to GTPB4 to maintain its conformation and coordinate removal of assembly factors during late nuclear maturation (Fig. 1G).

Pre-ribosome assembly requires the coordinated action of hundreds of assembly factors for the maturation of the small (SSU) and large subunit (LSU) (*1*), giving rise to numerous assembly intermediates along both pathways. Starting from our consensus maps, 3D classification identified six SSU processome states (Fig. 2A) and 14 pre-60s states (Fig. 3A) in native HeLa cells (fig. S2 to S4). We fitted known molecular models into our maps to identify previously reported early-to-late assembly states and the locations of specific proteins, revealing, in addition, the presence of previously uncharacterized intermediates in human pre-ribosome assembly. We placed the distinct structural classes into temporal order via tracing of the binding and unbinding of certain ribosomal protein and assembly factors to reconstitute an *in situ* pre-ribosome assembly pathway, which we describe in the following sections (Fig. 2A and Fig. 3A).

**Fig. 2.**
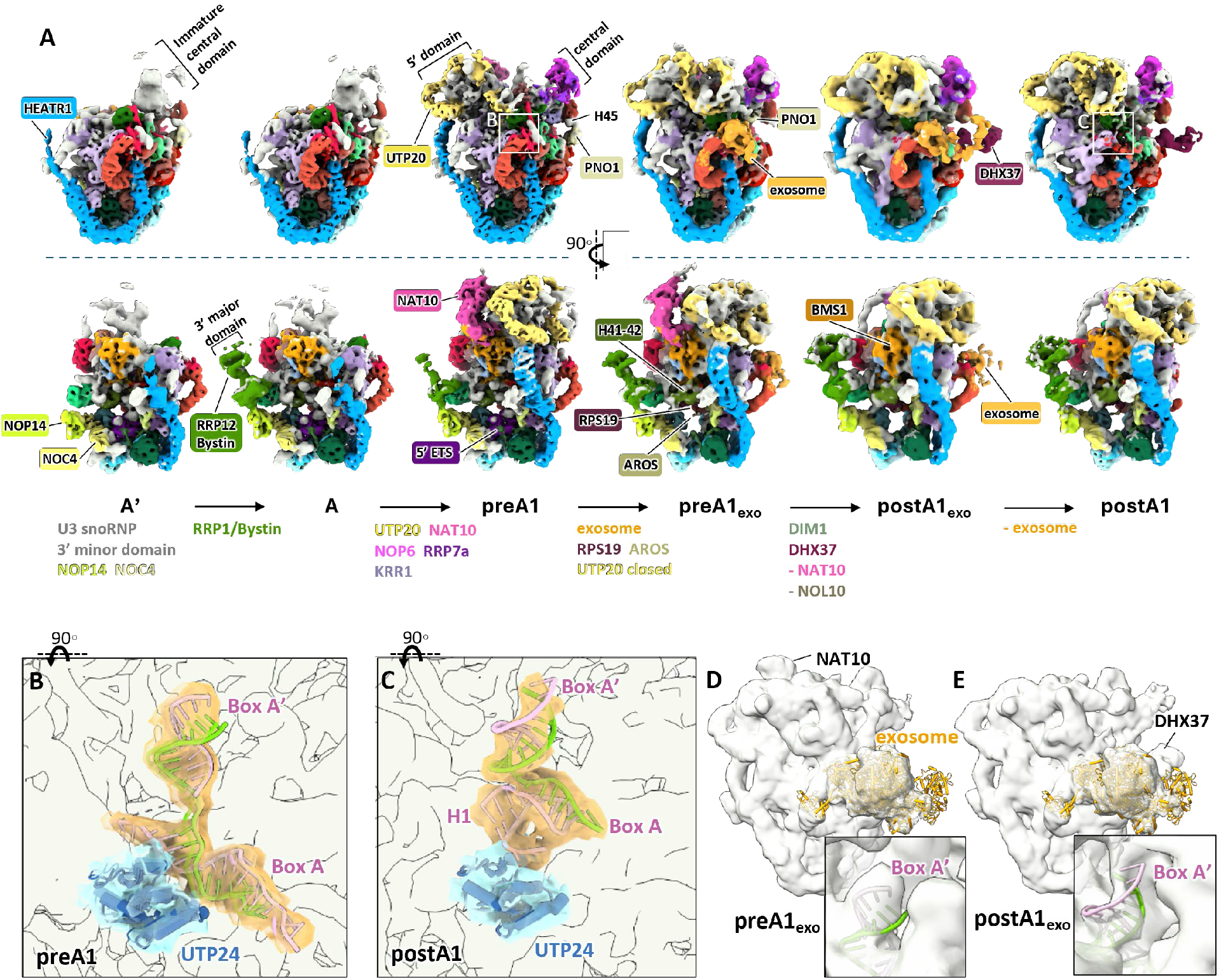
Assembly of the human SSU processome in the nucleolus. (**A**) Cryo-ET maps of distinct structural states of the human SSU processome, arranged in a putative temporal order of nucleolar assembly. The direction of the assembly pathway is indicated by the arrows. Assembly factors and modules are colored and labeled and conformational changes are indicated. Known molecular models (PDB 7mq8 and 7mqa) were fitted into the cryo-ET maps as detailed in the Methods. (**B** and **C**) Conformational changes of box A and box A’ before and after A1 cleavage catalyzed by endonuclease UTP24 (zoom in regions are indicated by white boxes in (A)). Dismantling of 18S rRNA, U3 snoRNA from PDB 7mq8 for state pre-A1 and from PDB 7mqa for state postA1. (**D** and **E**) The human exosome complex (PDB 9y8n) fitted into the SSU processome states preA1_exo_ and post A1_exo_ map. Zoom ins highlight the outward conformational change of box A’ after A1 cleavage.

**Fig. 3.**
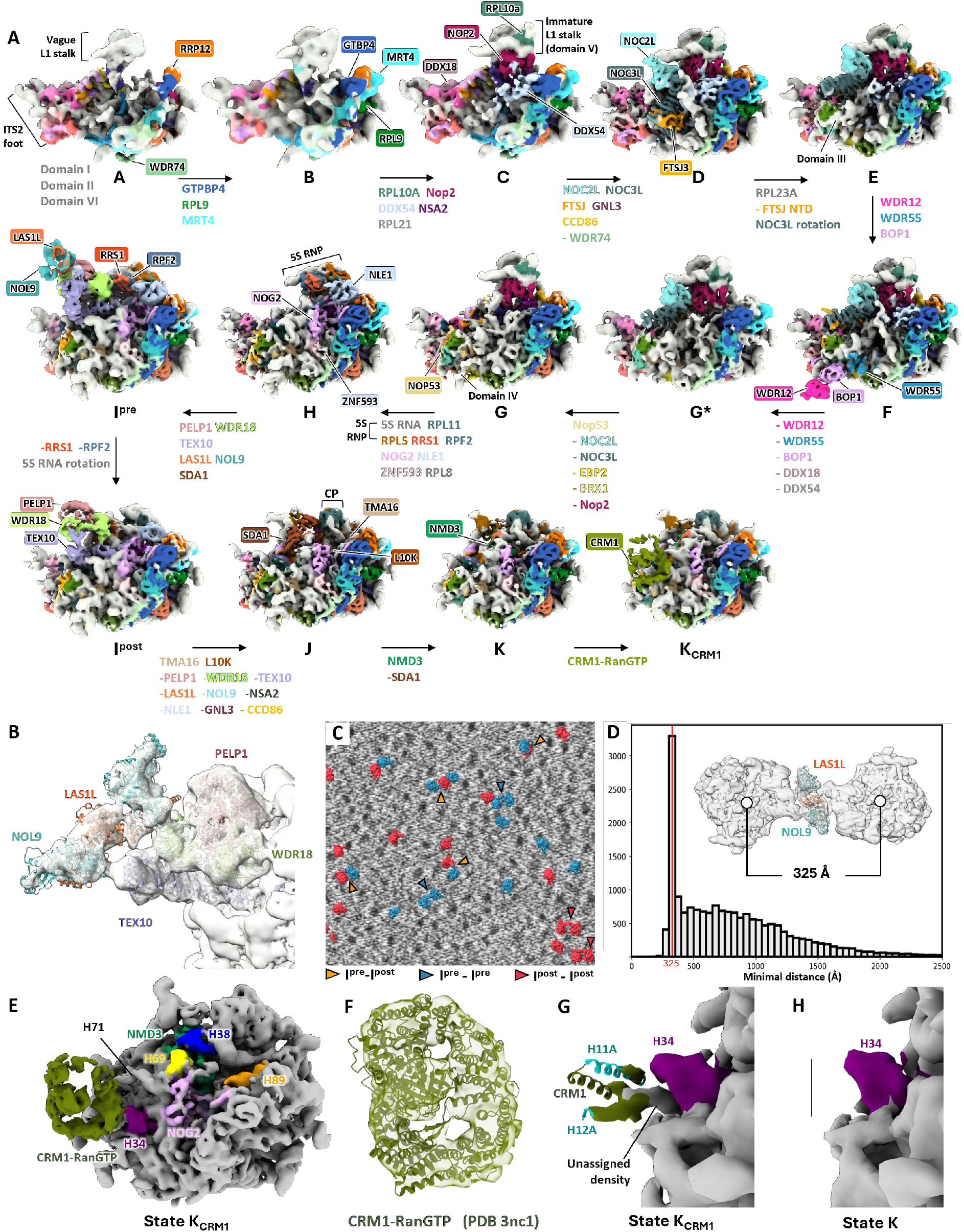
Assembly of the human pre-60S in the nucleolus. (**A**) Cryo-ET maps of distinct structural states of the human pre-60S, arranged in suggested temporal order of assembly in the nucleolus. Known molecular models (8fkp, 8fkr, 8fkt, 8fkv, 8fkx, 8fky, 8fkz, 8fl0, 8fl2, 8fl6, 8fla, 8flc, 8idt, 8ir3, 6lsr, 6lu8, 8fhr, 9yun, 3nc1) were fitted into the cryo-ET maps as detailed in the Methods. CP: central protuberance. (**B**) The human LAS1L-NOL9 complex model (PDB 26lk) and RIX1 complex (PDB 8fl2) fitted in pre-60S state I^pre^. (**C**) Distribution of pre-60S state I^pre^ and I^post^ in a representative tomogram of a native HeLa cell nucleolus. Pre-60S dimers indicated by arrowheads. Colors indicate different combinations of I^pre^ and I^post^ states in dimers. (**D**) Distance distribution between rixosome containing pre-60S particles. A distinct peak suggests an average distance of 325Å between pre-60S particles in dimers. (**E**) EM map of CRM1-RanGTP bound to pre-60S in presence of NMD3. **(F**) Mouse CRM1-RanGTP (PDB 3nc1) locally fitted into the cryo-ET density of pre-60S state K_CRM1_. (**G** and **H**) Unassigned helical density in state K_CRM1_ in proximity of the NES-binding groove between HEAT repeat helices H11A and H12A in state K_CRM1_. This density is missing in state K.

### Stepwise assembly of SSU processome in the human nucleolus

The human SSU processome is a giant precursor of the 40S ribosomal subunit, which couples RNA folding to subsequent RNA cleavage and processing (*18*). Assembly factors for the SSU processome are reported to organize into biochemically stable subcomplexes with 18S rRNA and U3 snoRNA. These include UTPA and UTPB complexes that chaperone the nascent 5’-ETS, U3 snoRNA that binds to 5’-ETS particles and serves as a platform for constructing further subdomains, the Mpp10 complex which associates with the 3’ major domain, the UTPC complex located on the central domain, and the Bms1-Rcl1 heterodimer that binds to the 5’ domain (*13, 21*). In line with reports in *Saccharomyces cerevisiae* and *Chaetomium thermophilum*, our in-cell structures of the SSU processome show a similar reverse order of co-transcriptional 18S rRNA subdomain integration (Fig. 2A); here, although the transcription of the 18S rRNA occurs from the 5’ end to the 3’ end, large parts of the 18S 3’ and central domains assemble first into the SSU processome before the 5’ domain is integrated. We characterized two new states represented by the first two SSU processome classes in the series, state A’ and state A, containing assembled U3 snoRNP, 3’ minor domain and an immature UTPC-chaperoned central domain. The NOP14-NOC4 module is already integrated in both states A’ and A, but the RRP12-Bystin module is missing in state A’. This indicates that the 3’ major domain is exposed which makes it available for pseudouridylation by box H/ACA snoRNA snR83 (*22, 23*). The identification of states A′ and A suggests an early stepwise assembly pathway of the human SSU processome, in which the exposure of the 3′ major domain in state A′ likely permits early snoRNP-guided rRNA modifications before the compaction of the rRNA and the masking of target sites by assembly factors.

The following preA1 state resembles the well-known SSU processome particle, in which the 5’ domain is incorporated together with UTP20 and the Kre33/NAT10 module, and is the last state proceeding A1 cleavage reported for human SSU processome intermediates (*12, 13, 24*) (Fig. 2A, state preA1). The next reported state after A1 cleavage, postA1, is also identified in our data and is characterized by the departure of the Kre33/NAT10 module, compaction of UTP20, repositioning of PNO1, and the recruitment of DHX37, DIM1 and RPS19 with its chaperon AROS (*13*) (Fig. 2A, state postA1). Cleavage at the A1 site between 5’-ETS and 18S rRNA releases the 5’ end of the 18S rRNA and allows the formation of RNA helix H1. At the same time, the box A’ heteroduplex formed by the 18S rRNA central domain and U3 snoRNA move upwards serving as evidence for A1 cleavage (*12*) (Fig. 2, B and C). In addition to the previously reported preA1 and postA1 human SSU processome states and the newly identified A′ and A states, we identified two further intermediates, preA1_exo_ and postA1_exo_, which are characterized by the presence of additional RNA exosome density as detailed below.

### Visualization of RNA exosome engagement during A1 cleavage

By visualizing the pre-ribosome assembly pathway in the cellular context, our *in situ* approach allowed transient interactions to be captured that can be difficult to preserve *ex situ*. Accordingly, we resolved binding of the RNA exosome, which is responsible for 5’-ETS degradation (*25, 26*), to both preA1 and postA1 particles in our data (Fig. 2A, states preA1_exo_ and postA1_exo_). Although similar structures are reported in yeast (*27*), and partial density of the exosome is resolved in the human SSU processome state postA1 (*28*), how the exosome engages the human SSU processome remained unclear.

While the resolution of the RNA exosome density in our data was relatively low because of its flexibility, it was sufficient to confidently fit the model of the human RNA exosome-Mtr4 complex (*29*) (Fig. 2, D and E). Compared to the preA1 state, preA1_exo_ particles show UTP20 in a closed conformation. The conformational change of UTP20 offers a binding site for the AROS N-terminal tail and recruits RPS19 (*13*). The incorporation of RPS19/AROS stabilizes RNA helix H41 of 18S rRNA and induces formation of H42 (Fig. 2A). The PNO1 relocates to the interior together with RNA helix H45 which brings the 5’ end of 18S rRNA closer to nuclease UTP24 for later A1 cleavage. The 5’-ETS density is absent in the preA1_exo_ particles (Fig. 2A) because it is dislodged and relocated by the docking of the exosome onto the UTP6 platform (*13, 28*). The 18S rRNA box A’ undergoes an upwards conformational change from state preA1_exo_ to postA1_exo_ (Fig. 2, D and E), suggesting complete A1 cleavage while the exosome remains bound. These states provide new mechanistic insight into how the exosome facilitates 18S rRNA processing.

### Maturation of the human pre-60S in the human nucleolus

Our data also visualized the pre-60S assembly pathway within the nucleolus (Fig. 3A). A hallmark of human pre-60S assembly is the early establishment of organizational hubs that coordinate pre-rRNA folding and recruit assembly factors for successive maturation steps (*9*). The early assembly precursors identified in our data closely resemble structures reported *in vitro*, confirming the consistency of our results with earlier studies. Assembly begins with the formation of rRNA root helices in state A, followed by the recruitment of GTPB4 in state B to initiate the formation of the peptidyl transferase center (PTC). In state C, RPL10A and the NIP7/NOP2 complex are incorporated, stabilizing the L1 stalk in an immature conformation. DDX54 joins at this stage, remodels rRNA and acts as a platform for recruitment of the FTSJ3-NOC2L-NOC3L complex in state D (*9*). In our data, a weak density for the L1 stalk was already visible in states A and B. The NOC2L-NOC3L complex rearranges from a downward orientation in state D to an upward one in state E, enabling formation of domain III. In state F, BOP1, WDR12, and WDR55 chaperone the maturation of domain III (*9*). We identified a new intermediate state G*, which shows subsequent removal of helicases DDX18 and DDX54, followed by the release of NOP2, the NOC2L-NOC3L complex, EBP2, and BRX1 in state G. This allows for the rotation of the L1 stalk and creates space for incorporation of the non-rotated 5S RNP after NOG2 tethering in state H.

We also observed additional late maturation intermediates prior to nuclear export. It is known that following 5S RNP binding and rotation, the rixosome, consisting of the RIX1 subcomplex (PELP1-WDR18-TEX10), and the LAS1L-NOL9 complex (*30*), is recruited via SDA1 to catalyze ITS2 cleavage (*9*). In the newly identified state I^pre^, the 5S RNP remains unrotated while the rixosome is bound, whereas in state I^post^, the LAS1L-NOL9 density becomes weaker concurrent with the rotation of 5S RNP, indicating increased flexibility of the complex. These observations suggest that the rotation of 5S RNP is not strictly coupled to rixosome binding. We discuss these new states in further detail below. The central protuberance (CP) forms in state J upon detachment of NLE1, CCD86, and GNL3 and incorporation of TMA16 and RPL10. Finally, SDA1 is replaced by 60S ribosomal subunit export adaptor NMD3 in state K, which binds RNA helix H38 in an extended conformation. We additionally resolved a density in a new state K_CRM1_ that could be explained by the nuclear export receptor CRM1-RanGTP when fitting the model into the density. The early recruitment of the CRM1-RanGTP complex suggests that pre-ribosome maturation in the nucleolus has already progressed to a late maturation stage poised for subsequent nuclear export. In summary, we recapitulated eight structures similar to previous studies (A, B, C, D, E, F, G, H) (*9*–*11*), while extending the pre-60S assembly pathway by six new states (state G*, I^pre^, I ^post^, J, K, K_CRM1_).

### Cryo-ET reveals the intact human rixosome and its bridging of pre-60S dimers

One essential irreversible step of 28S rRNA maturation is the removal of ITS2 catalyzed by the rixosome (*30*). Within the rixosome complex, LAS1L provides endoribonuclease activity required for pre-rRNA cleavage (*31*), while NOL9 functions as a 5’-OH polynucleotide kinase that phosphorylates the cleavage products (*32*), thereby enabling subsequent trimming by 5’-3’ exoribonucleases. Although structures of the RIX1 subcomplex and the LAS1L-NOL9 subcomplex have been determined separately by cryo-EM (*30, 33*), the structure of the fully assembled rixosome remained unknown. Our *in situ* analysis revealed the architecture of the intact human rixosome on the pre-60S particles and provides insight into how the catalytic LAS1L-NOL9 module is positioned to mediate ITS2 cleavage on pre-60S ribosomes (Fig. 3A, state I^pre^ and state I^post^).

The LAS1L-NOL9 module in the rixosome bound to state I^pre^ showed a dimer of LAS1L-NOL9 heterodimers (Fig. 3, A and B) arranged in C2 symmetry, forming a heterotetramer with two independent catalytic centers (Fig. 3, A and D), which is consistent with the structure of the reconstituted LAS1L-NOL9 complex (*30, 33*). Fitting atomic models of the human LAS1L-NOL9 complex and the RIX1 complex into the cryo-EM density of state I^pre^ demonstrates that the LAS1L-NOL9 complex directly interacts with TEX10 and WDR18, thereby bridging the two submodules into a unified complex (Fig. 3B). Analysis of the 3D spatial distribution of rixosome-bound particles revealed a substantial population of pre-60S dimers bridged by shared LAS1L-NOL9 heterotetramers. These pre-60S dimers comprise I^pre^–I^pre^ homodimers, I^pre^–I^post^ heterodimers, and a smaller fraction of I^post^–I^post^ homodimers (Fig. 3C), suggesting that rixosome-mediated dimerization persists across successive maturation states. Consistent with this observation, the distribution of minimal interparticle distances exhibited a pronounced peak at approximately 325 Å (Fig. 3D), corresponding to the characteristic separation within these dimeric assemblies (fig. S6, A and B). In the I^post^ state, the RIX1 complex underwent a ∼9° rotation, concomitant with the rotation of the 5S RNP (fig. S6, C and D). As a result, the relative orientation between particles differed across dimer classes: I^pre^–I^pre^ homodimers displayed the smallest interparticle angle (27◦), followed by I^pre^–I^post^ heterodimers (35◦), and finally I^post^–I^post^ homodimers (43◦) (fig. S6E). The formation of pre-60S dimers bridged by the LAS1L-NOL9 complex may represent a mechanism to simultaneously exploit both catalytic sites of the LAS1L-NOL9 heterotetramer, potentially enhancing the efficiency or coordination of ITS2 processing during late nucleolar maturation of the pre-60S subunit.

### Early nucleolar recruitment of the nuclear export machinery

One unexpected observation in our data was the presence of a CRM1-RanGTP complex bound to the pre-60S particles in the nucleolus (Fig. 3, E and F, state K_CRM1_). The nuclear export adaptor NMD3 contains a nuclear export signal (NES) at its C-terminus and is considered essential for recruiting CRM1 during pre-60S nuclear export (*34, 35*). In nuclear pore-associated pre-60S particles in yeast, CRM1-RanGTP localizes near the L1 stalk in proximity to the NMD3 NES (*36*). In contrast, in our newly resolved state K_CRM1_, CRM1 occupies a distinct position adjacent to helix H34 in domain III of the 28S rRNA, far from both the L1 stalk and the NES of NMD3 (Fig. 3E).

Despite this distinct CRM1 position, the overall architecture of state K_CRM1_ closely resembles that of yeast nuclear pore-associated pre-60S particles (*36*). Consistent with previous reports that NMD3 associates with pre-60S particles at late stages of assembly (*37, 38*), we observed both NMD3 and NOG2 in our newly resolved state K_CRM1_ (Fig. 3E). Here, NMD3 binds to RNA helix H38 and maintains it in an open conformation, while central helices H69 and H71 remain immature because steric hindrance by NOG2 prevents their conformational flipping. Compared with the yeast pre-60S particles, state K_CRM1_ lacks the yeast-specific export factors MEX67, MTR2, ARX1, and ECM1, but contains human L10K (fig. S7).

Compared with state K, state K_CRM1_ displays an additional rod-like density. This suggests that a helical peptide binds CRM1 between HEAT repeat helices H11A and H12A (Fig. 3, G and H), which form the canonical NES-binding cleft (*39*). This density likely represents a stabilized NES from another so far unidentified ribosomal protein or biogenesis factor that is distinct from NMD3 (Fig. 3G). Our observations reveal a new CRM1-RanGTP binding site on pre-60S particles that is distinct from that observed in nuclear pore-associated pre-60S intermediates. This suggests that pre-60S particles may harbor multiple CRM1 interaction sites, potentially contributing to the high nuclear export efficiency of pre-ribosomes in live cells (*40*), and that CRM1 can adopt different binding modes depending on the subcellular context, inside and outside the nuclear pore.

Importantly, the identification of state K_CRM1_ within the nucleolus implies that CRM1 may interact with the pre-60S particles earlier than previously thought, potentially mediating the release of pre-60S particles from the nucleolus to the nucleoplasm. Supporting this model, inhibition of CRM1 by leptomycin B, which covalently blocks the NES-binding cleft, leads to the nucleolar accumulation of 28S rRNA (*41*), indicating that the interaction with CRM1 is required for efficient nucleolar exit of pre-60S.

### Pol I inhibition impacts the release of pre-60S from the nucleolus

Pol I transcription governs the structural integrity of the nucleolus. Inhibition of Pol I activity is known to reorganize nucleoli towards more spherical shapes and to induce the formation of distinct structures containing Pol I, called nucleolar caps (*42*) (fig. S8A). To investigate how inhibition of Pol I transcription and such rearrangements of the nucleolus affect the pre-ribosome maturation process, we treated HeLa cells with low-dose ActD to selectively inhibit Pol I transcription (*43*) (fig. S8A, Methods) and applied our CLEM-guided cryo-ET workflow to target both the nucleolus and nucleolar caps regions (fig. S9).

Following a similar subtomogram analysis approach described above, we mapped all instances of each pre-ribosome state back to the tomograms of the native and ActD perturbed nucleoli to generate 3D particle maps. In the untreated cells, both SSU processome and pre-60S particles predominantly localized within the GC area, defined by NPM1 fluorescence (Fig. 4A). Notably, all pre-60S states, including K_CRM1_, were found within the nucleolus. This suggests that late maturation can occur here, while subsequent steps presumably only comprise transport through the nucleoplasm and nuclear pore mediated by CRM1 (15). Also notably, we did not observe any significant maturation gradient of pre-ribosome intermediates across the human nucleolus.

**Fig. 4.**
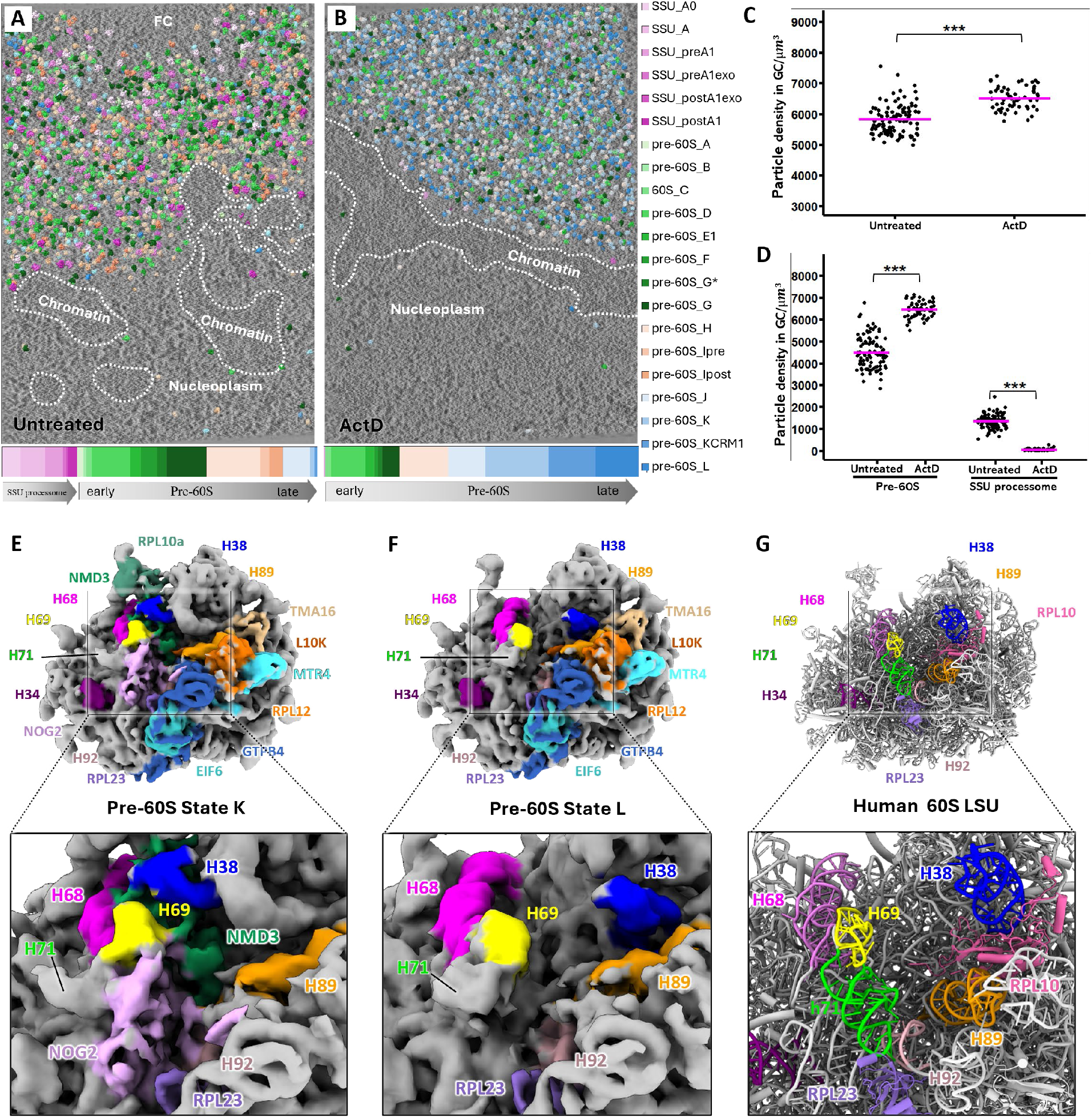
Pol I inhibition alters pre-ribosome assembly. (**A** and **B**) Particle map of pre-ribosomes in a representative untreated HeLa cell and in a representative HeLa cell treated with 5 nM ActD for 3 hours. The bottom bars show the proportion of different states of SSU processome and pre-60S following a temporal order of assembly from the left to the right. (**C**) Density of total pre-ribosomal particles in the GC before and after ActD treatment. Asterisks denote significance levels determined using the t-test (***P < 0.001). (**D**) Density of pre-60S and SSU processome in the GC before and after ActD treatment. (**E** and **F**) Cryo-ET maps of pre-60S states K and L, relevant ribosome biogenesis factors and rRNA helices colored according to published molecular models (PDB 8idt, PDB 8qyx, and PDB 6lsr). (**G**) Molecular model of the cytosolic human 60S ribosomal subunit (PDB 8qyx), showing mature conformations of rRNA helices H38, H68, H69, H71, and PTC helix H89 and H92.

The spatial distribution of ActD-treated cells revealed a similar predominant nucleolar localization of pre-ribosomal particles (Fig. 4, A and B, fig. S8, B to D). However, nucleolar particle concentration increased significantly upon ActD treatment, from mean ∼5,800 particles/µm^3^ in untreated to mean ∼6,500 particles/µm^3^ in ActD-treated cells (Fig. 4C). The small- and large-subunit precursors exhibited opposite changes in abundance upon Pol I inhibition. SSU processome particles were almost entirely absent from the nucleolus, while the total pre-60S particle concentration increased (Fig. 4D). In the pre-60S population, early states A to G showed a decreased proportion. Rixosome-bound pre-60S particles (state I^pre^ and state I^post^) were absent, indicating stalled ITS2 cleavage. Late states K and K_CRM1_ showed a dramatic increase, indicating stalled release of mature particles from the nucleolus (Fig. 4, A and B). Together, these changes suggest that Pol I inhibition suppresses early ribosome biogenesis while promoting accumulation of late, mature pre-60S particles within the nucleolus, consistent with impaired nucleolar exit.

Given that Pol I inhibition caused accumulation of late pre-60S particles in the nucleolus (Fig. 4B), we asked whether this reflected a general defect in nuclear export. However, ActD treatment had minimal effects on global nuclear export (fig. S10, A and B). We therefore assessed nuclear CRM1 levels after ActD treatment to test for a nucleolus-specific transport defect. CRM1 showed low nucleolar signal before ActD treatment, but accumulated after treatment (fig. S10C), consistent with previous reports (*41, 44*). Nucleolar CRM1 levels began to increase 40 min after ActD treatment and continued to rise in the following hour (fig. S10D). These results indicate that pre-60S accumulation in the nucleolus is specific to a defect in CRM1-mediated pre-60S transfer from the nucleolus to the nucleoplasm rather than a general nuclear export defect.

Interestingly, manual segmentation of perinucleolar heterochromatin indicated chromatin reorganization upon ActD treatment (Fig. 4, A and B, and fig. S9). In native HeLa cells, chromatin formed discontinuous regions at the GC surface that may allow pre-ribosome transit into the nucleoplasm (Fig. 4A, and fig. S9). Upon ActD treatment, chromatin reorganized into a continuous layer around the NPM1-labelled GC (Fig. 4B, and fig. S9), consistent with an increase in DNA fluorescence intensity surrounding the GC (fig. S10A). It is possible that the continuous layer of chromatin forms a physical barrier that restricts pre-60S release.

In addition, we resolved a previously uncharacterized pre-mature state L in the ActD-treated nucleoli (Fig. 4F). Compared with state K, NOG2 and NMD3 were absent in state L. The central helices H69 and H71 were flipped towards H92, adopting a near-mature conformation following NOG2 departure (*45*). The 28S rRNA helix H38 adopted a closed, mature-like conformation following NMD3 dissociation. However, GTPB4 remained bound in state L, maintaining H89 in an immature, open conformation (Fig. 4, E, F, G). State L is reminiscent of the previously reported pre-60S state K3 derived from *ex situ* data (*9*) (fig. S11), and structurally represents an intermediate, pre-mature conformation positioned between late nucleolar pre-60S particles (Fig. 4E) and the cytosolic 60S subunit (Fig. 4G). This state possibly corresponds to a reversible intermediate, arising from the dynamic association of nucleolar assembly factors in the absence of cytosolic maturation factors.

## Conclusion

Our study illuminates human pre-ribosome maturation under near-native conditions in intact nucleoli. Using CLEM-guided cryo-ET and subtomogram analysis, we captured known and novel structural intermediates of both the SSU processome and pre-60S at secondary-structure level details despite the crowded nucleolar environment. The *in situ* analysis presented here uniquely preserves transient and labile interactions. This enabled the visualization of the RNA exosome contacting human SSU processome intermediates during early pre-rRNA processing. We also identified a fully assembled rixosome bound to pre-60S particles and revealed pre-60S dimerization mediated by the LAS1L–NOL9 heterotetramer. In addition, we identified CRM1 association with pre-60S particles within the nucleolus at a non-canonical binding site near rRNA helix H34, which may serve to monitor 28S rRNA maturation and facilitate licensing of nucleolar exit.

By tracing individual particles back to their precise cellular locations, we provide evidence that most assembly steps occur within the nucleolus, including those of pre-60S states I^pre^ to K_CRM1_, whereas pre-60S maturation steps following 5S RNP incorporation were thought to take place in the nucleoplasm (*9, 10*). While previous work reported a spatial unidirectional gradient of pre-ribosomal particles across their maturation steps in nucleoli of *C. reinhardtii*, we did not observe a comparable gradient in our HeLa nucleoli data. In contrast to *C. reinhardtii*, human nucleoli contain multiple FC/DFC units that function as independent ribosome biogenesis initiation sites (*4, 14, 42*). It is possible that the presence of multiple, spatially distributed assembly centers generates overlapping fluxes of pre-ribosomal intermediates, leading to mixing within the nucleolar environment and thereby limiting the appearance of a clear unidirectional gradient.

While Pol I transcription is a major factor in determining nucleolar structure, its impact on the pre-ribosome maturation process has remained unclear. We show that inhibition of Pol I transcription results in depletion of the SSU processome in the nucleolus, while at the same time leads to accumulation of late-stage pre-60S particles. This differential response highlights uncoupling of stress-induced regulations of SSU and LSU assembly. The enrichment of late pre-60S particles suggests stalling of pre-ribosome assembly and impaired nucleolar exit. This may originate from altered GC material properties (*46*–*49*) and/or the formation of a chromatin barrier. Indeed, we show that Pol I inhibition leads to reorganization of perinucleolar chromatin and the formation of a chromatin barrier on the GC surface. In contrast, the rapid SSU turnover (*6*) may allow SSU precursors to escape the nucleolus before establishment of these alterations.

In conclusion, our study provides an integrated perspective wherein pre-ribosome assembly is not only governed by sequential factor recruitment of different ribosomal components and assembly factors but is also shaped by spatial organization and cellular states. This work thus provides a framework for understanding how cells modulate ribosome production in response to changing physiological or stress conditions.

## Acknowledgments

We acknowledge support by the EMBL Advanced Light Microscopy Core Facility and access and support by the EMBL Imaging Centre. X.Z. acknowledges support from the EMBL International PhD programme. We thank J. Bartho, S. Unger, Z. Yang for support in cryo-EM data acquisition, T. Hoffmann and EMBL IT support for computational and data storage support, members of the Cuylen-Häring, Mahamid and Müller groups for support and discussion, and J. Cheng for sharing the model of the full-length human LAS1L-NOL9 complex.

## Author contributions

Conceptualization: S.C.-H., J.M., C.W.M.

Methodology: X.Z., Y.H., H.K.H.F., S.C.-H., J.M.

Investigation: X.Z., Y.H., H.K.H.F.

Visualization: X.Z., Y.H.

Funding acquisition: S.C.-H., J.M., C.W.M.

Project administration: C.W.M.

Supervision: S.C.-H., J.M., C.W.M.

Writing – original draft: X.Z.

Writing – review & editing: Y.H., H.K.H.F., S.C.-H., J.M., C.W.M.

## Data, code, and materials availability

Cryo-ET maps will be deposited in the Electron Microscopy Data. Accession numbers will be provided with publication.

## Supplementary Materials

Materials and Methods

Supplementary Text

Figs. S1 to S11

Table S1

References (*51*–*76*)

