## Supplementary Data for "In-cell structures visualize human pre-ribosome assembly in the nucleolus"

**The PDF file includes:**

Materials and Methods
Supplementary Text
Figs. S1 to S11
Tables S1
References

### Materials and Methods

#### Cell culture

All cell lines used in this study were regularly tested and confirmed to be free of mycoplasma contamination. Cell line sources and authentication are summarized in the Reagents and Tools table. All experiments were performed in HeLa-Kyoto cells described previously (*51*). Cells were maintained in Dulbecco's modified Eagle medium (DMEM; Thermo Fisher Scientific) supplemented with 10% (v/v) fetal bovine serum (FBS; Thermo Fisher Scientific), 1% (v/v) penicillin-streptomycin (Sigma Aldrich), and 1 mM sodium pyruvate (Thermo Fisher Scientific), hereafter referred to as WT medium. For live-cell imaging, cells were cultured in FluoroBrite DMEM (Thermo Fisher Scientific) supplemented with 10% (v/v) FBS, 1% (v/v) penicillin-streptomycin, 1 mM sodium pyruvate, and 1% (v/v) GlutaMAX (Thermo Fisher Scientific).

#### Generation of stable cell lines

Stable cell lines expressing SNAP-NPM1, FBL-mEGFP, mEGFP-NPM1, FBL-SNAP or H2B-EBFP2 were generated by lentiviral transduction. Lentiviral particles were produced using the corresponding lentiviral transfer plasmids together with a VSV-G envelope plasmid (Addgene #12259) and the pCMV-R8.74 packaging plasmid (Addgene #22036). Transduced cells were selected for 2 weeks in WT medium supplemented with either 6 µg/mL blasticidin S hydrochloride (Sigma-Aldrich) or 300 µg/mL hygromycin B (Thermo Fisher Scientific). Following selection, fluorescent cells were isolated by either bulk sorting or single-cell sorting into 24-well plates or 96-well plates using a FACSAria™ Fusion flow cytometer (BD Biosciences). Sorted cell populations or clones were assessed by cell morphology and fluorescence microscopy to confirm the expected protein localization.

For endogenous N-terminal tagging, donor repair plasmids were designed to insert the Halo-tag at the N-terminus of UBF or the SNAP-tag at the N-terminus of RPA194. Donor sequences were synthesised and cloned into pUC57 (Biomatik), and contained approximately 700 bp homology arms flanking the UBF or RPA194 start codon, with the corresponding tag sequence inserted between the homology arms. Endogenous knock-in cell lines were generated using either conventional CRISPR/Cas9-mediated genome editing or a double Cas9 nickase strategy. For the Halo-tag knock-in at the UBF N-terminus, cells were electroporated with two Cas9 nickase–sgRNA plasmids and a Halo-tag donor plasmid using the Neon Transfection System. After 1 week, cells were labelled with 50 nM Halo-TMR and isolated into 96-well plates by flow cytometry. Clones were screened by genotyping PCR, immunoblotting and fluorescence microscopy to confirm successful knock-in and the expected subcellular localization. For the SNAP-tag knock-in at the RPA194 N-terminus, cells were electroporated with a single plasmid expressing wildtype Cas9 and sgRNA and a SNAP-tag donor plasmid using the Neon Transfection System. After 1 week, cells were labelled with 100 nM SNAP-SiR and isolated into 96-well plates by flow cytometry. Clones were screened by genotyping PCR and fluorescence microscopy to confirm successful knock-in and the expected subcellular localization.

#### Cryo-ET sample preparation

R 1.2/20 SiO2 grids (Au 200 mesh, Quantifoil) were plasma cleaned using a Fishione M1070 plasma cleaner (2 min; Ar/$O_2$ 90:10), and micropatterned with 30-µm fibronectin circles in the center of grid squares, as described (*52*).

Micropatterned grids were placed in a 35 mm cell culture dish (Greiner) with 1 ml DMEM (Thermo Fisher Scientific). Cells from a 25T flask at ~80% confluency were trypsinized and resuspended into 1 ml DMEM (Thermo Fisher Scientific), filtered into a test tube with cell strainer snap cap 35 µm mesh (Corning), and added to the dish. The adherence state of cells on the grids was confirmed after 45 min incubation, and grids were transferred into a new dish with fresh medium. Before plunge freezing, cells were treated with 200 µM oleic acid (Sigma) together with 0.2 µM BODIPY 558/568 (Sigma) for 45 min to induce lipid droplet formation used for accurate registration of the post-milling cryo-fluorescence and cryo-TEM lamella maps. Detailed below. Two triple labeled HeLa cell lines were used for cryo-ET data acquisition: a cell line with mEGFP-NMP1, Halo-UBF, and SNAP-RPA194 was stained with 0.2 µM JFX Halo-646 (Janelia Materials) to localize Pol I; a cell line with FBL-mEGFP, Halo-UBF, and SNAP-NPM1 (*53*) was stained with 0.2 µM JFX Halo-549 and JFX SNAP-650 (Janelia Materials) for 15 min to visualize all three nucleolar components. For ActD treatment, cells were additionally incubated in 5 mM ActD (Sigma) for 3 h before plunge freezing. For vitrification, grids were blotted from the backside for 4 sec using a Leica EM GP2 plunge freezer at 65% humidity, 37°C, and plunged into liquid ethane. Grids were stored in sealed grid boxes in liquid nitrogen.

The cells were subsequently milled using an Aquilos FIB-SEM (Thermo Fisher Scientific) on a shuttle with a 45◦ pretilt angle. Before milling, grids were sputter coated with metallic platinum (1 kV, 10 mA, 10 Pa, 30 sec), followed by organometallic platinum coating via the gas injection system (GIS) with a thickness of 0.5 to 1 µm. Using AutoTEM software, cells were milled at a stage tilt of 9◦ while stepwise decreasing the gallium ion beam current (1 nA, 0.5 nA, 0.3 nA, 50 pA and 30 pA), aiming at lamellae thickness of 150 nm. A final manual polishing step was applied, at a stage tilt at 10◦ and with a current of 30 pA. A final sputter-coating with platinum (1 kV, 10 mA, 10 Pa, 4 sec) was applied to the polished lamellae to reduce charging. Two out of eight cryo-ET datasets (described below) were acquired on lamellae generated with fluorescence-targeted milling: fluorescence stacks of cells were acquired with a wide-field fluorescence light microscope (iFLM) equipped with a 20x/0.7 NA Zeiss Epiplan-Apochromat objective, integrated in the Aquilos2 column. The fluorescence stacks were registered on the SEM grid maps using Maps 3.8 (Thermo Fisher Scientific) and 3D targeting was done with AutoTEM software. Grids used in these experiments were plunge frozen together with 1 µm FluoSpheres polystyrene orange (540/560) beads (Invitrogen) as fiducials for fluorescence registration and 3D targeting.

<u>On-lamella CLEM</u>

Polished lamellae were loaded on to a Zeiss LSM900 cryo-Airyscan2 confocal microscope. Overview images were acquired using a 5x air objective in widefield mode to localize lamellae. Z-stacks with 0.5 µm spacing were acquired in Airyscan mode with 488 nm, 546 nm, and 640 nm lasers lines using a 100x/0.75 NA air objective at low laser power (less than 5%). In addition, reflection mode images were acquired for each z-plane. Z-stacks were 2D Airyscan processed using the ZEN Blue software. Subsequently lamella maps were acquired on a Titan Krios G3i (Thermo Fisher Scientific) at a magnification of 6,500x (pixel size 14.02 Å, defocus of -90 µm) and stitched using SerialEM or IMOD etomo BlendMontage (*54*). The sum projections of the fluorescence stacks were registered on to the TEM lamella maps using the lipid droplets as fiducials in ImageJ BigWarp and using the Affine mode (*55*).

<u>Cryo-ET data acquisition</u>

Data were acquired on a Titan Krios G3i (Thermo Fisher Scientific) transmission electron microscope operated at 300 keV equipped with post-column energy filter (Gatan) and K3 direct electron detector (Gatan) using SerialEM software (*56*). Tilt series were collected from -60◦ to +60◦ at 3◦ increment using the dose-symmetric scheme (*57*). Total fluence was ~140 $e^-/Å^2$, pixel size was 3.425 Å at nominal magnification 26,000x, defocus range was set from -3 to -7 µm. Targeted nucleolar areas were selected based on the CLEM results described above. In total, 212 tilt series were collected in native HeLa cells and 93 in ActD treated cells. To generate initial references for template matching (described below), 27 untreated tomograms were acquired using a Volta phase plate (VPP) (*58*).

Tomogram reconstruction, particle picking and subtomogram analysis

Tilt series were motion- and CTF corrected in WarpTools (*59*) or Warp v1.09 (*60*) (for VPP data), aligned with AreTomo2 (*61*), and reimported into WarpTools or Warp for tomogram reconstruction. Tomograms were reconstructed at 13.70 Å/pixel for visualization and particle picking.

For the small VPP dataset of untreated cells, a human SSU processome state preA1 map (EMD 23936) was low-pass filtered to 30 Å and used as the reference for 2D template matching using SUSAN v1.0 (*62*, *63*). Detected particles, together with particles manually picked in EMAN2 (*64*), were extracted at a pixel size of 5 Å in Warp, followed by 3D global classification in RELION v4.0.1 (*65*). Global classification identified 2 classes of pre-ribosomal particles: SSU processome and pre-60S. Subsets of SSU processome and pre-60S were selected after classification for RELION 3D auto-refine (fig. S2, maps ① and ②). The refined maps were low-pass filtered to 50 Å and used as initial references for 3D template matching.

For the defocus datasets, 3D template matching of SSU processome and pre-60S were performed using these maps in PyTom-GPU (*66*, *67*) with an angular search of 7.5° and a spherical mask. Template matching peaks within the lamella masks, 1500 for each tomogram, were filtered and extracted. Datasets collected from untreated and ActD treated cells were combined to obtain consensus maps for the pre-ribosomes. Subtomograms were reconstructed at 5 Å/pixel using WarpTools and initially classified in RELION using 3D global classification with a T value of 0.2 or 0.4. Duplicates were removed with a RELION function (overlap criterion: <150 Å) and particles from each class were refined in RELION. The resulting averages (fig. S2, maps ⑦- ⑪) were used as references for multi-species M of geometrical deformations in the tilt series (*59*, *68*). Another round of M-correction was performed separately by combining classes into SSU processome and pre-60S species to improve the resolution of the consensus maps.

Tomograms were reconstructed at 13.70Å/pixel using the M-refined alignments, and a new round of template matching was performed using the same parameters described above. Due to the high heterogeneity of pre-60S particles, newly refined maps (fig. S3, map ⑨- ⑪) representing distinct assembly states were used as references, which was important to complete the particle lists and to reduce bias toward late-state pre-60S particles, which dominated the consensus map. The expected particle numbers were estimated according to the numbers of meaningful particles retained in each tomogram in the last round of classification. The template matching peaks from these three references (fig. S3, map ⑨- ⑪) were combined and duplicates were removed. Pre-60S particles from untreated and ActD treated cells were processed separately here. For the first global classification, each job was replicated three times, the good particles from these three jobs were combined and duplicates were removed. After several rounds of global classification with a T value of 0.2 or 0.4, followed by classification with a T value of 2 or 4 (fig. S3) without alignment and premature refinement, classified particles of each group were selected, duplicate particles removed, and subsequently refined in RELION.

Focused classification was then performed with a T value of 2 or 4, but without alignments (fig. S4). Masks for focused classification were generated by volume rendering surrounding the fitted molecular models of biogenesis factors (RRP12-NOP14 and NAT10 from PDB 7mq8, DHX37 from 7mqa, exosome from PDB 9y8n, NOP2 and MRT4 from PDB 8fkt, NOC2L/NOC3L from PDB 8fkv, NOC2L/NOC3L rotated from PDB 8fkx, WDR12/BOP1 from PDB 8fky, rixosome from PDB 8fl2 and 26lk, CRM1-RanGTP from PDB 3nc1). In order to reduce variance, classifications were repeated 3-5 times, and the most consistent set of particles was selected. The output classes were subjected to additional classification and a round of local refinement.

Once classified, particles in the same states from untreated and ActD treated cells were merged for local refinement in RELION and subsequent multi-species M refinement (*59*, *65*, *68*). Output half-maps from M were re-imported into RELION for post-processing, where Fourier Shell Correlation (FSC) curves and final resolutions were calculated for all identified complexes (fig. S5). Masks for post-processing were generated using denoised maps from M with 6-pixel binary and 8-pixel soft expansion.

Model fitting and visualization

Ribosomal proteins, biogenesis factors and RNA modules in published molecular models (PDB 7mq8, 7mqa, 8fkp, 8fkr, 8fkt, 8fkv, 8fkx, 8fky, 8fkz, 8fl0, 8fl2, 8fl6, 8fla, 8flc, 8idt, 8ir3, 6lsr, 6lu8, 8fhr, 9yun, 3nc1, 26lk) were extracted separately and used for rigid-body fitting into the pre-ribosome maps in ChimeraX 1.10 (*69*), to identify individual components in the complexes. The cryo-ET maps were then colored according to the fitted molecular models for visualization. Visualization of local resolution was performed in ChimeraX by coloring the EM maps with the local resolution files of the refined maps from M.

Pre-ribosome states analysis in individual tomograms

After classification and refinement, all pre-ribosomes and their states were mapped back to their original tomograms using the ChimeraX plugin ArtiaX (*69*, *70*). The nucleolar pre-ribosome concentrations were calculated using 99 tomograms from untreated cells and 52 tomograms from ActD treated cells. The nucleolar volume in each tomogram was first defined by high particle-density clusters using DCSCAN (*71*) and then verified using the fluorescence signals.

Pre-60S dimer analysis

The particle coordinates and orientations after RELION refinement were analyzed using custom Python scripts. In brief, particles were grouped by tomogram, and pairwise Euclidean distances between their centers were computed in three dimensions. For each particle, the minimal distance to its nearest neighbor was determined. For rixosome bound pre-60S particles (states $I^{pre}$ and $I^{post}$), distance distributions and summary statistics were generated across all tomograms, and results were exported and plotted in histograms (Fig. 3D), the most abundant peak was detected at 325Å. The same distance analysis was performed for all pre-60S states. Distances were calculated both within particle classes and between classes by combining datasets and selecting pairs belonging to specified states. For each state or state pair, the fraction of dimers with inter-particle distances within a target range (325 ± 25 Å) was calculated. These fractions were assembled into a matrix summarizing enrichment across all group combinations and visualized as a heatmap (fig. S6A). For the distance-orientation analysis, particle orientations were converted into unit vectors by applying Euler rotations (ZXZ convention) to a reference axis. Pairwise relationships were computed within each tomogram using a k-dimensional-tree-based neighbor search (*72*) with a maximum distance cutoff (5000 Å). For each particle pair, the Euclidean distance and relative orientation were calculated, where the orientation angle was defined as the angle between the corresponding unit vectors. The distance–orientation distributions were computed as two-dimensional histograms (fig. S6B), smoothed, and used to identify dominant structural populations. Orientation distributions were analyzed independently to identify preferred angular relationships (fig. S6E).

Nuclear transport reporter assay

A reporter plasmid was synthesized to express emiRFP670 fused to a nuclear export signal (NES) and mEGFP fused to the myc nuclear localization signal (mycNLS). A self-cleaving T2A peptide sequence was inserted between emiRFP670-NES and mEGFP-mycNLS to enable expression of both reporters from a single plasmid.

Cells stably expressing SNAP-NPM1 and H2B-EBFP2 were seeded in 35 mm confocal dish (LabTek) and immediately transfected with 20 ng/µl plasmid encoding reporter gene cassette (emiRFP670-NES-T2A-mEGFP-mycNLS) using jetPrime (Polyplus). H2B-EBFP2, an H2B fusion with enhanced blue fluorescent protein 2, was used to visualise chromatin. Two days later, cells were treated with 5 nM Act D or 5 ng/ml Leptomycin B (LMB) for 3h and then fixed with 4% formaldehyde. Single focal planes were acquired on a Zeiss LSM780 confocal microscope with a Plan-Apochromat 63×/1.4 NA Oil DIC M27 operated using ZEN Black software.

For quantification of reporter enrichment in subcellular regions, images were analyzed using CellProfiler (*73*). Nuclei were segmented from the DNA signal using the adaptive Otsu method (*74*). To define the cytoplasm, nuclear masks were expanded by 5 and 20 pixels. The 5-pixel-expanded masks were then subtracted from the 20-pixel-expanded masks to generate ring-shaped perinuclear regions, which were used as cytoplasmic masks. Mean reporter fluorescence intensities were then measured in each of the subcellular region. The output from CellProfiler was further analyzed in R for data visualization and statistical analysis. Non-transfected cells were excluded on the basis of a manually assigned threshold for cytoplasmic emiRFP670-NES intensity. Nuclear enrichment of the reporter signal was calculated as the ratio of nuclear to cytoplasmic signal intensity. For statistical analysis, data normality and homogeneity of variance were assessed using the Shapiro–Wilk test and Levene's test, respectively. Statistical comparisons were performed using the Kruskal–Wallis test, followed by Dunn's post hoc test.

Live cell imaging

For the nucleolar cap formation, Cells stably expressing three nucleolar marker proteins (Halo-UBF, FBL-SNAP, and mEGFP-NPM1) were seeded in a Lab-Tek chambered slide (Thermo Fisher Scientific) one day before imaging. Halo-tag and SNAP-tag were labeled with 100 nM Halo-Tetramethylrhodamine (TMR; Promega) and 100 nM SNAP-silicon rhodamine (SiR; New England Biolabs), respectively, according to the manufacturer's instructions. To induce nucleolar cap formation, cells were treated with 5 nM ActD (Thermo Fisher Scientific) for 2 h. Images were acquired on an Olympus IXplore SpinSR microscope equipped with a UPLSAPO 100×/1.35 NA objective and an incubation chamber maintaining 37°C and 5% $CO_2$. Z-stacks spanning 3 µm were acquired at 0.28 µm intervals.

To assess CRM1 enrichment, an EGFP-CRM1 expression plasmid was generated using FLAG-hCRM1 (Addgene #17647) as the backbone. The FLAG tag was replaced with an EGFP fragment amplified from an EGFP-containing plasmid using Gibson assembly. For time-lapse imaging of CRM1 enrichment, cells stably expressing SNAP-NPM1 were seeded in a 35 mm confocal dish (LabTek) and immediately transfected with 20 ng/µl plasmid encoding EGFP-CRM1. Two days after transfection, SNAP-NPM1 and DNA were labeled with 100 nM SNAP-SiR and 1× SPY555-DNA (Spirochrome) for 1h, respectively. Cells were then washed and maintained in imaging medium. Imaging was performed on an Olympus IXplore SpinSR microscope equipped with a UPLSAPO 100×/1.35 NA silicon oil objective and an incubation chamber maintaining 37°C and 5% $CO_2$. Time-lapse z-stacks spanning 5.5 μm were acquired at 0.5 μm intervals every 10 min for 2h. After the first frame was acquired, ActD was added to the medium at a final concentration of 5 nM.

To analyze the time-lapse data, the DNA channel was first converted into maximum-intensity projections. Nuclei were then segmented and tracked over time on the basis of the projected DNA signal using TrackMate (*75*) in Fiji (*76*). Based on the tracking results, individual nuclei were cropped across the time series. For each cropped nucleus at each time point, a single z-plane was selected on the basis of mean NPM1 intensity, and the resulting single-slice time-lapse images were analyzed in CellProfiler. Nuclei and nucleoli were segmented from the DNA and NPM1 signals, respectively, using the adaptive Otsu method. Nucleoli were assigned to their parent nuclei. Nucleoplasm masks were generated by subtracting nucleolar masks expanded by 6 pixels (~0.4 μm) from nuclear masks. Mean fluorescence intensities were measured in each compartment at each time point. The CellProfiler output was further analyzed in R. CRM1 nucleolar enrichment was calculated as the ratio of nucleolar to nucleoplasmic signal intensity.

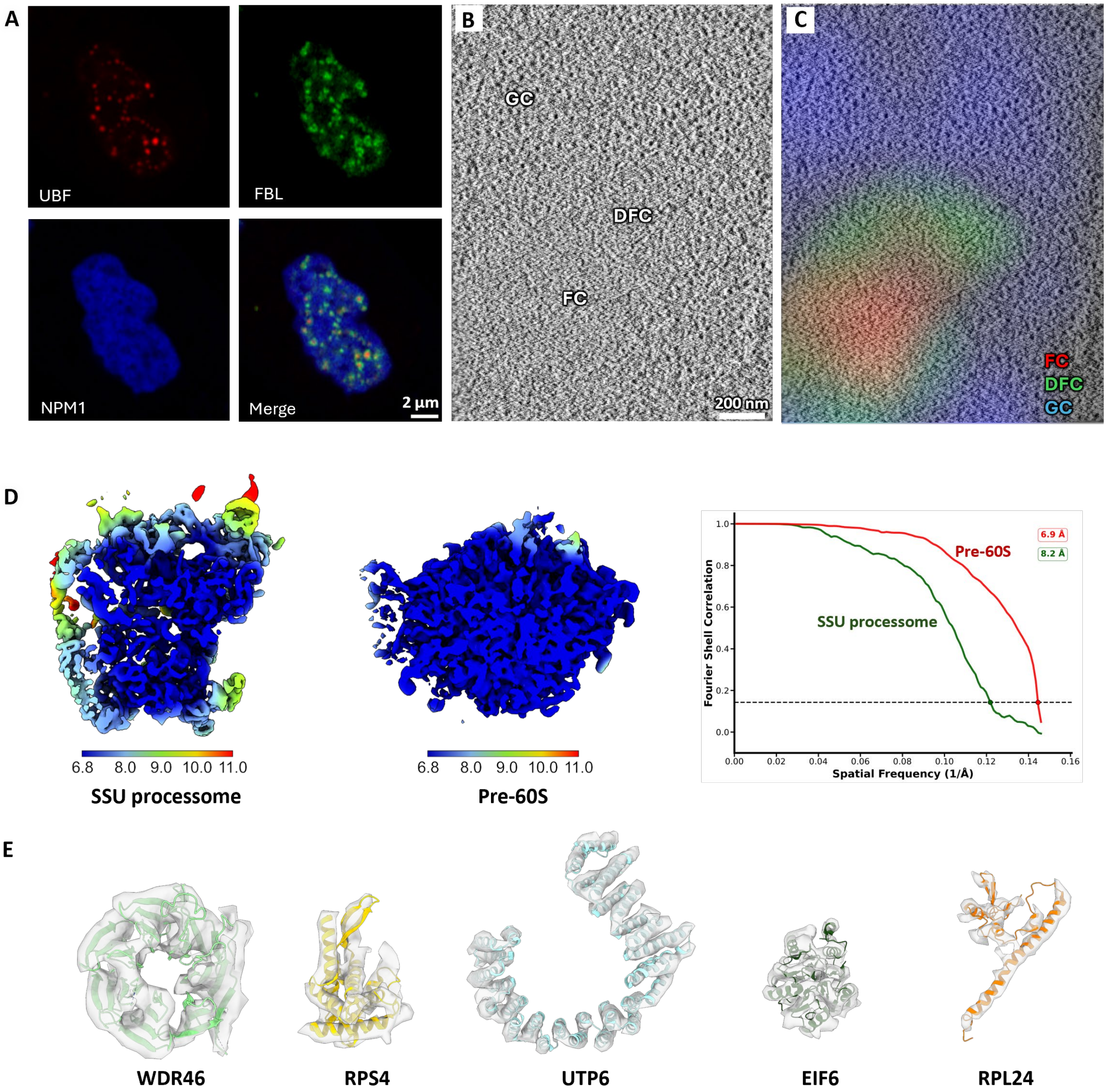


**Fig. S1. Human nucleoli and pre-ribosomes under cryo-electron microscope.**

(**A**) Architecture of a nucleolus in native HeLa cells. Live triple-labeled HeLa cell, FBL-mEGFP, Halo-UBF (JFX-Halo549), and SNAP-NPM1 (JFX-SNAP650), were imaged by Zeiss LSM880 Airyscan mode. (**B** and **C**) Central slice of a representative tomogram in a nucleolus in native triple-labeled HeLa cells. FC, DFC and GC are marked in (B) and correlated with fluorescence in (C). (**D**) Consensus cryo-EM maps of SSU processome and pre-60S particles. Fourier shell correlation (FSC) curves for SSU processome and pre-60S refinement, and the reported resolution value at FSC = 0.143. The Nyquist limit for the data is 6.85 Å. (**E**) Representative densities and models of ribosome biogenesis factors. Models for WDR46, RPS4, and UTP6 are from PDB 7mq8. Models for EIF6 and RPL24 are from PDB 8fla.

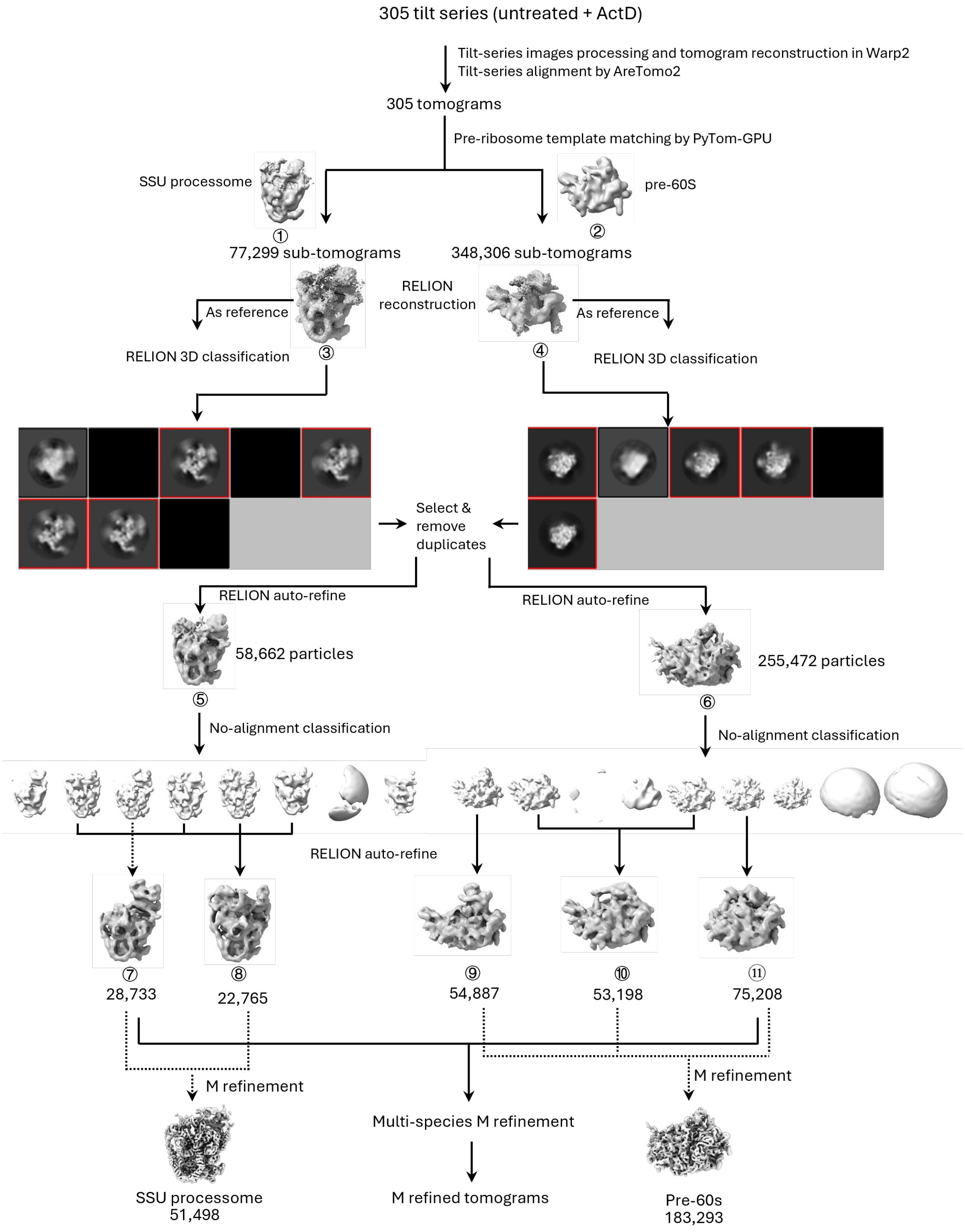


**Fig. S2. Initial classification and refinement of pre-ribosomes in untreated and ActD treated cells.**

A diagram of the image-processing workflow. For each reconstruction, a unique number identifier is assigned for tracking. The particles numbers are provided for each class.

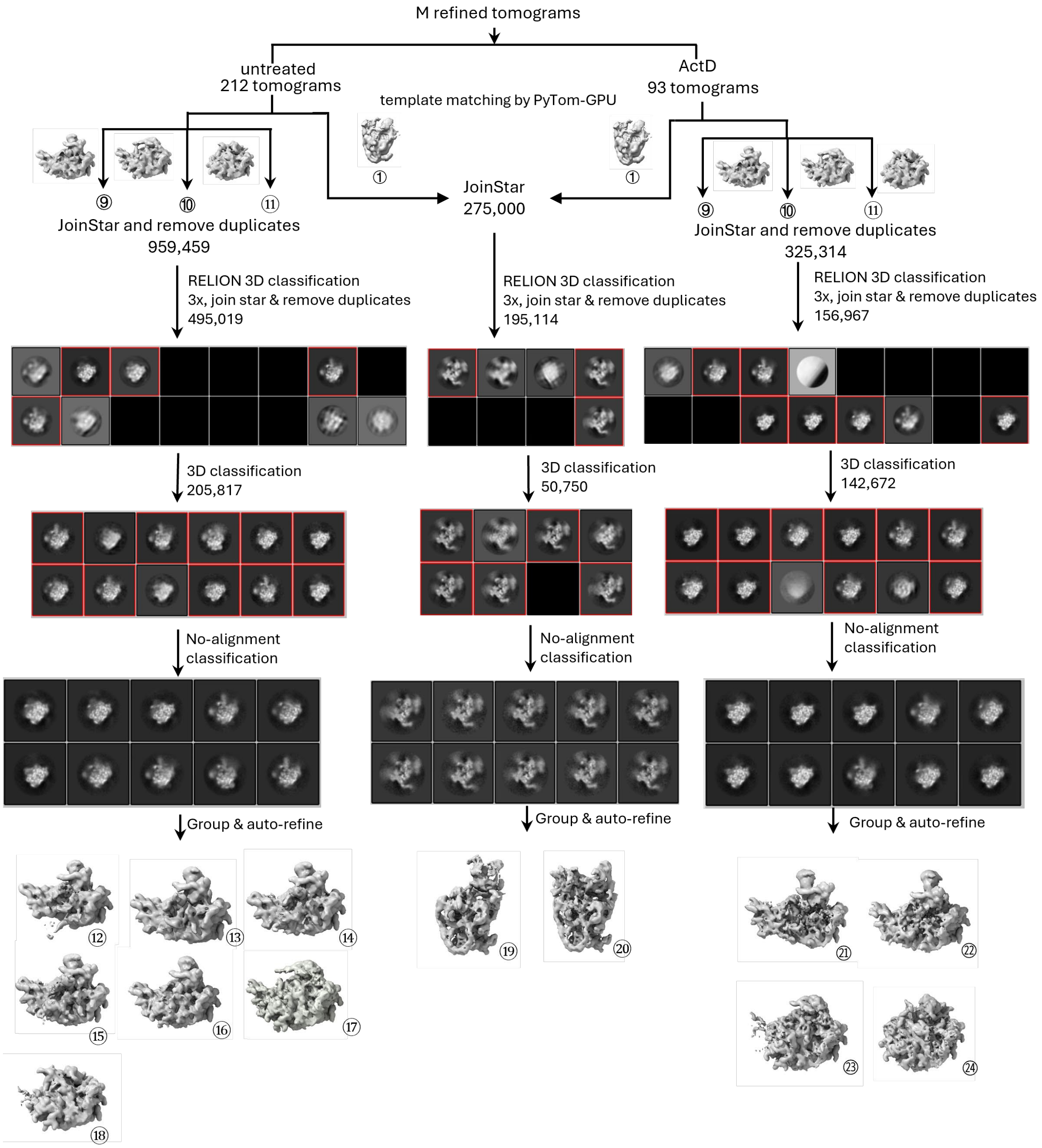


**Fig. S3. Global classification and refinement after M.**

Datasets from untreated and ActD treated cells are processed independently after one round of M refinement. 3D refinement was avoided before the last round of global classification to reduce overfitting. A unique number identifier is assigned for tracking for each reconstruct. The particles numbers are provided for each step.

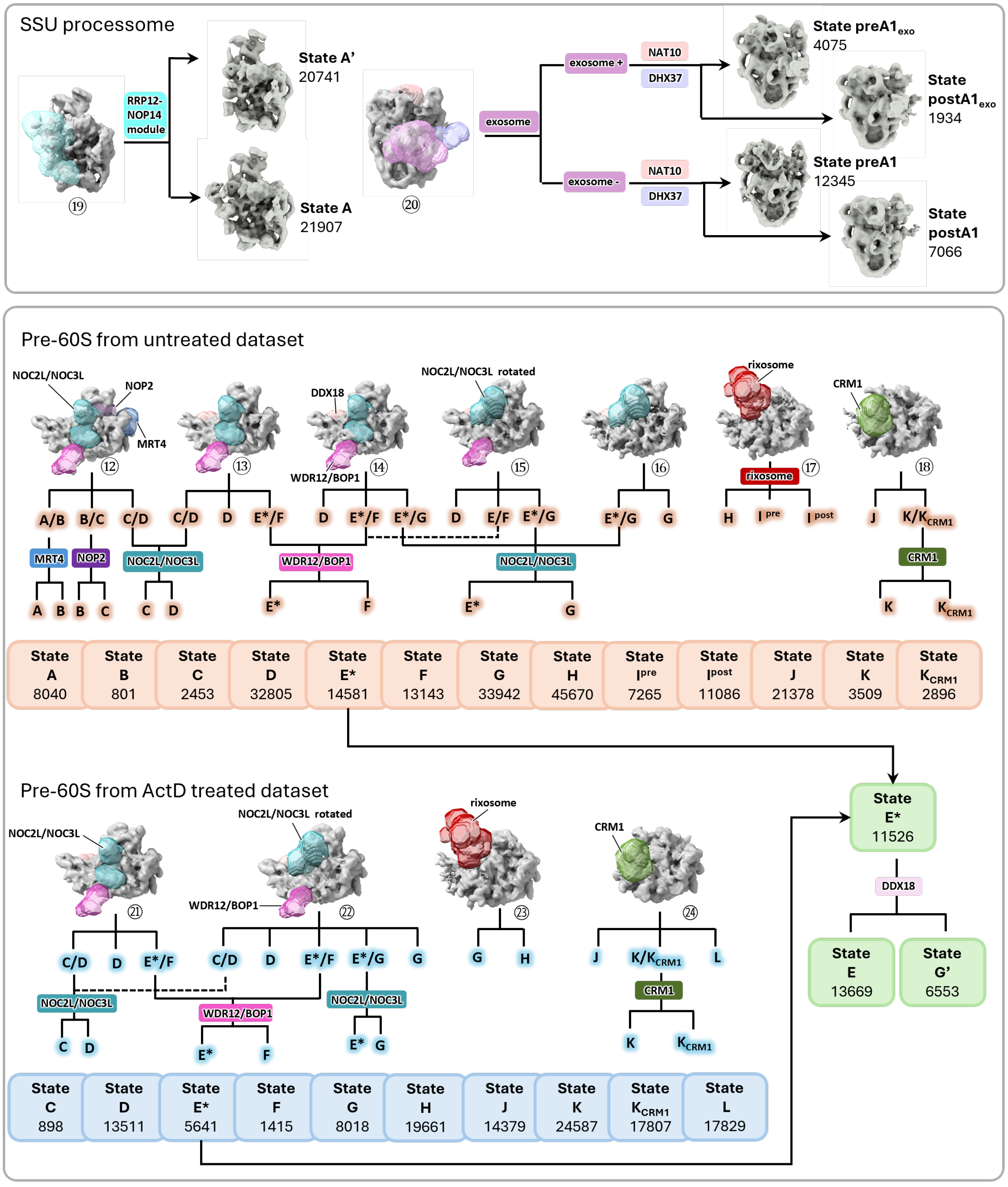


Same states of SSU processome and pre-60S from untreated and ActD treated datasets were merged

RELION 3D refinement & M refinement

**Fig. S4. Focused classification of pre-ribosomes.**

Multiple focused 3D classifications strategies used for the reconstruction of different biogenesis factors of the SSU processome and pre-60S. Masks used for the different classifications are color-coded and labeled. Molecular models used for mask generation: RRP12-NOP14 and NAT10 from PDB 7mq8, DHX37 from 7mqa, exosome from PDB 9y8n, NOP2 and MRT4 from PDB 8fkt, NOC2L/NOC3L from PDB 8fkv, NOC2L/NOC3L rotated from PDB 8fkx, WDR12/BOP1 from PDB 8fky, rixosome from PDB 8fl2 and 26lk, CRM1-RanGTP from PDB 3nc1. The same pre-ribosome states from untreated and ActD treated datasets are combined for M refinement to obtain better resolutions. The particles numbers are provided for each class.

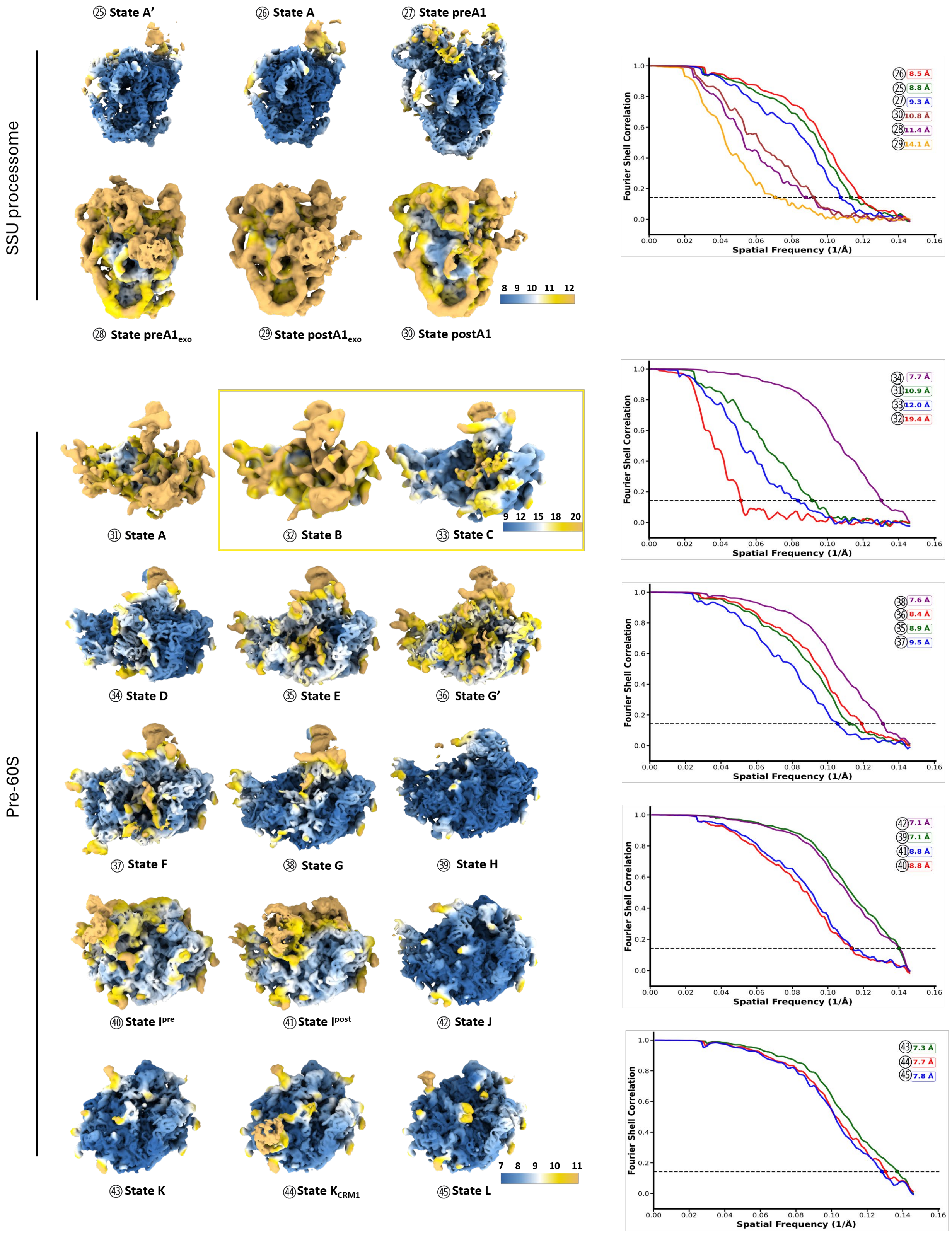
SSU processome
㉕ State A'
㉖ State A
㉗ State preA1
㉘ State preA1exo
㉙ State postA1exo
㉚ State postA1
8 9 10 11 12
Fourier Shell Correlation
Spatial Frequency (1/Å)
㉖ 8.5 Å
㉕ 8.8 Å
㉗ 9.3 Å
㉚ 10.8 Å
㉘ 11.4 Å
㉙ 14.1 Å
Pre-60S
㉛ State A
㉜ State B
㉝ State C
9 12 15 18 20
㉞ 7.7 Å
㉛ 10.9 Å
㉝ 12.0 Å
㉜ 19.4 Å
㉞ State D
㉟ State E
㊱ State G'
㊳ 7.6 Å
㊱ 8.4 Å
㉟ 8.9 Å
㊲ 9.5 Å
㊲ State F
㊳ State G
㊴ State H
㊵ State Ipre
㊶ State Ipost
㊷ State J
㊷ 7.1 Å
㊴ 7.1 Å
㊶ 8.8 Å
㊵ 8.8 Å
㊸ State K
㊹ State KCRM1
㊺ State L
7 8 9 10 11
㊸ 7.3 Å
㊹ 7.7 Å
㊺ 7.8 Å

**Fig. S5. Cryo-ET maps of pre-ribosome intermediates.**

Pre-ribosome maps colored by local resolutions. Fourier shell correlation (FSC) curves for SSU processome and pre-60S refinement, and the reported resolution value at FSC = 0.143. The Nyquist limit for the data is 6.85 Å.

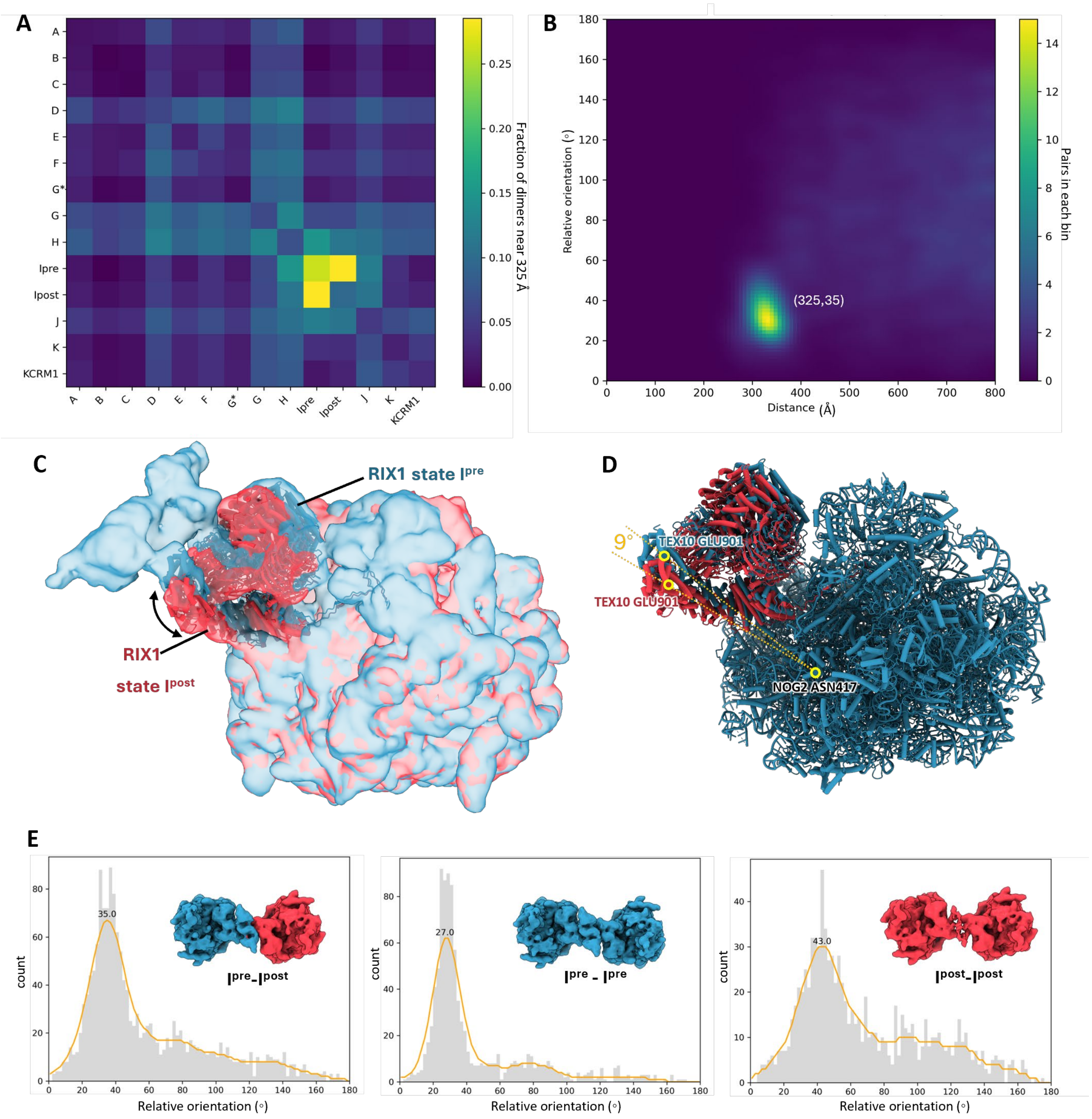


**Fig. S6. Dimerization of rixosome-bound pre-60S particles**

(**A**) Fractions of dimers with inter-particle distances within range 325 ± 25 Å across all pre-60S states. (**B**) Distance-orientation distribution of pre-60S state $I^{pre}$ combined with state $I^{post}$. Enrichment peak was marked out. (**C**) EM maps of pre-60S state Ipre and state Ipost with RIX1 complex model fitted in (PDB 8fl2). (**D**) Rotation of RIX1 complex was measured by the angle between atom pair NOG2 ASN417- TEX10 GLU901 in state $I^{pre}$ and state $I^{post}$. (**E**) Orientation distributions of particles in state $I^{pre}$ and state $I^{post}$ analyzed independently to identify preferred angular relationships. Pre-60S dimers were shown as displayed in 3D particle maps in ChimeraX.

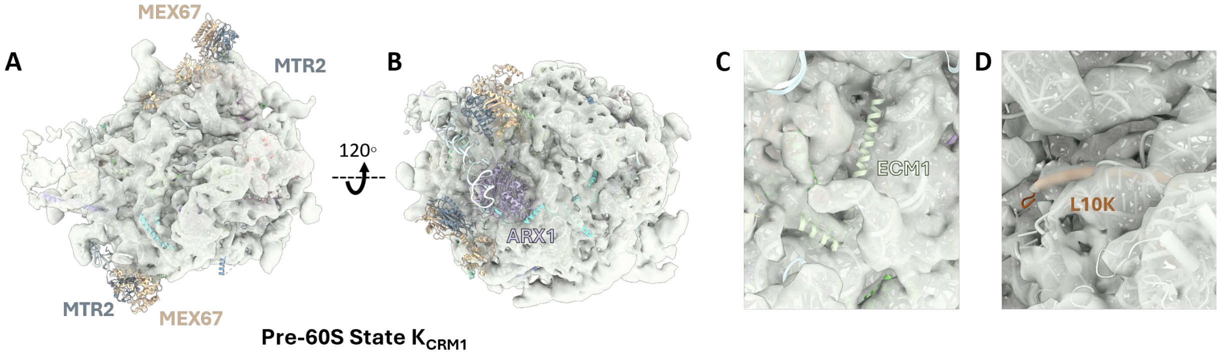


**Fig. S7. Comparison of pre-60S state $K_{CRM1}$ and yeast pre-60S trapped in NPC.**
Molecular model of yeast pre-60S trapped in NPC (PDB 8hfr) fitted into the EM map of pre-60S state $K_{CRM1}$.

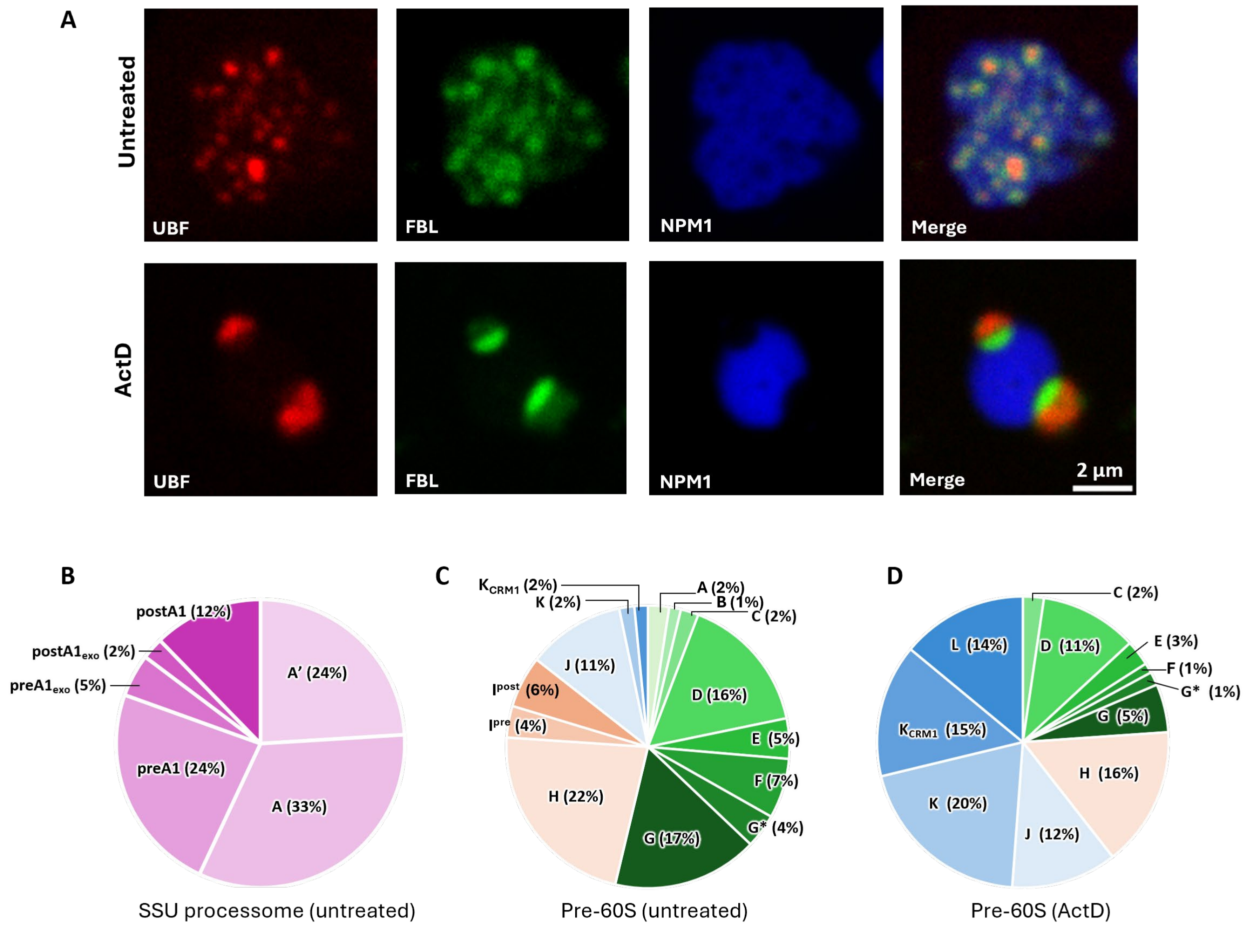


**Fig. S8. Proportion of the different states of SSU processome and pre-60S particles.**
(**A**) Formation of nucleolar cap induced by ActD treatment. Live triple-labeled HeLa cell, FBL-SNAP (SiR dye labelling), Halo-UBF (TMR dye labelling), and mEGFP-NPM1, were treated with 5 nM ActD for 3h and imaged by Olympus iXplore SPIN SR with UPLSAPO100XS. (**B**) Proportion of the different states of the SSU processome in untreated HeLa cells. (**C**) Proportion of the different states of pre-60S particles in the GC of untreated HeLa cells. (**D**) Proportion of the different states of pre-60S particles in the GC of ActD treated HeLa cells.

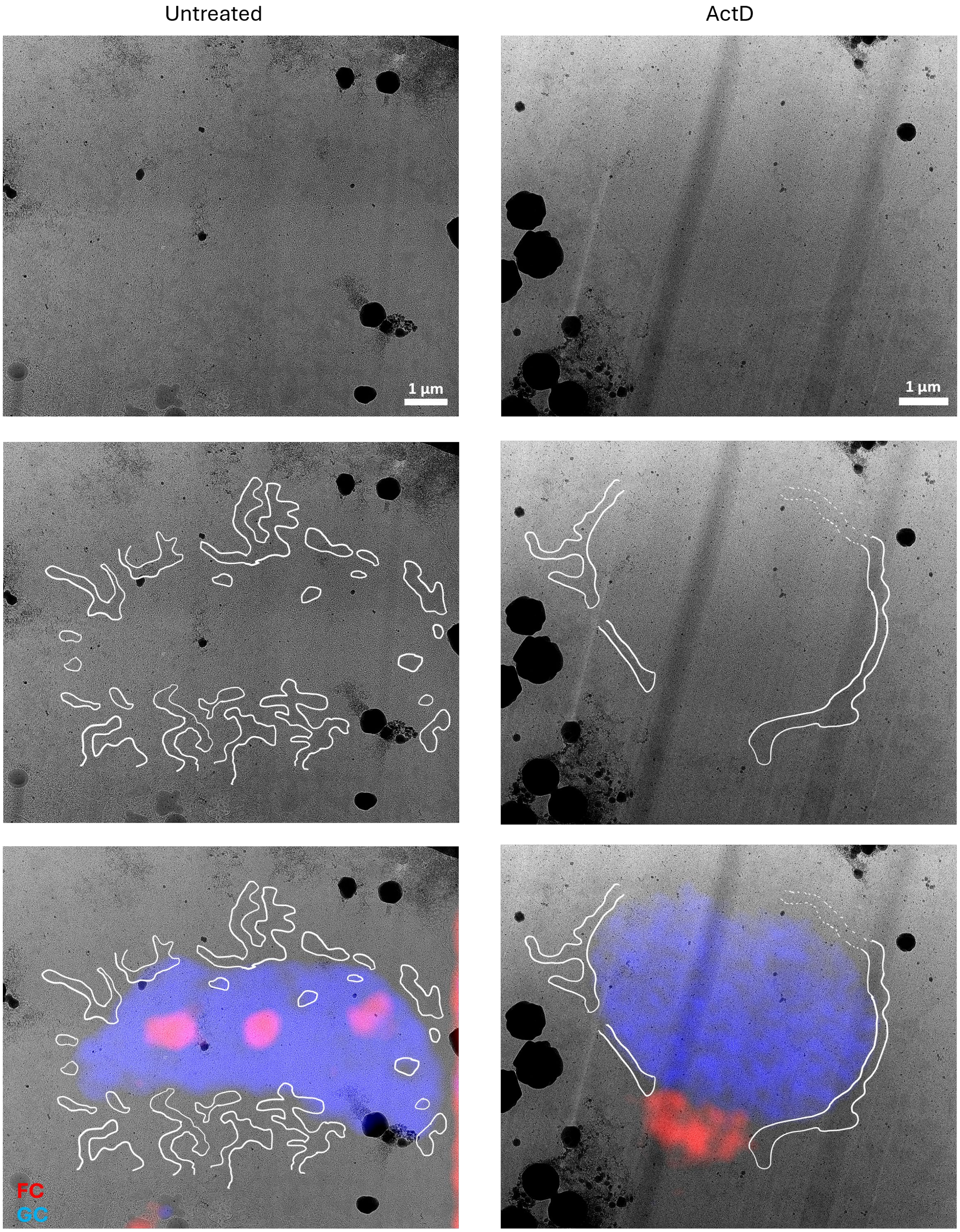


**Fig. S9. Lamella maps of nucleolus in TEM.**

Lamella maps of nucleolus in untreated and ActD treated HeLa cells. Perinucleolar heterochromatin was segmented manually and overlayed with the fluorescence image.

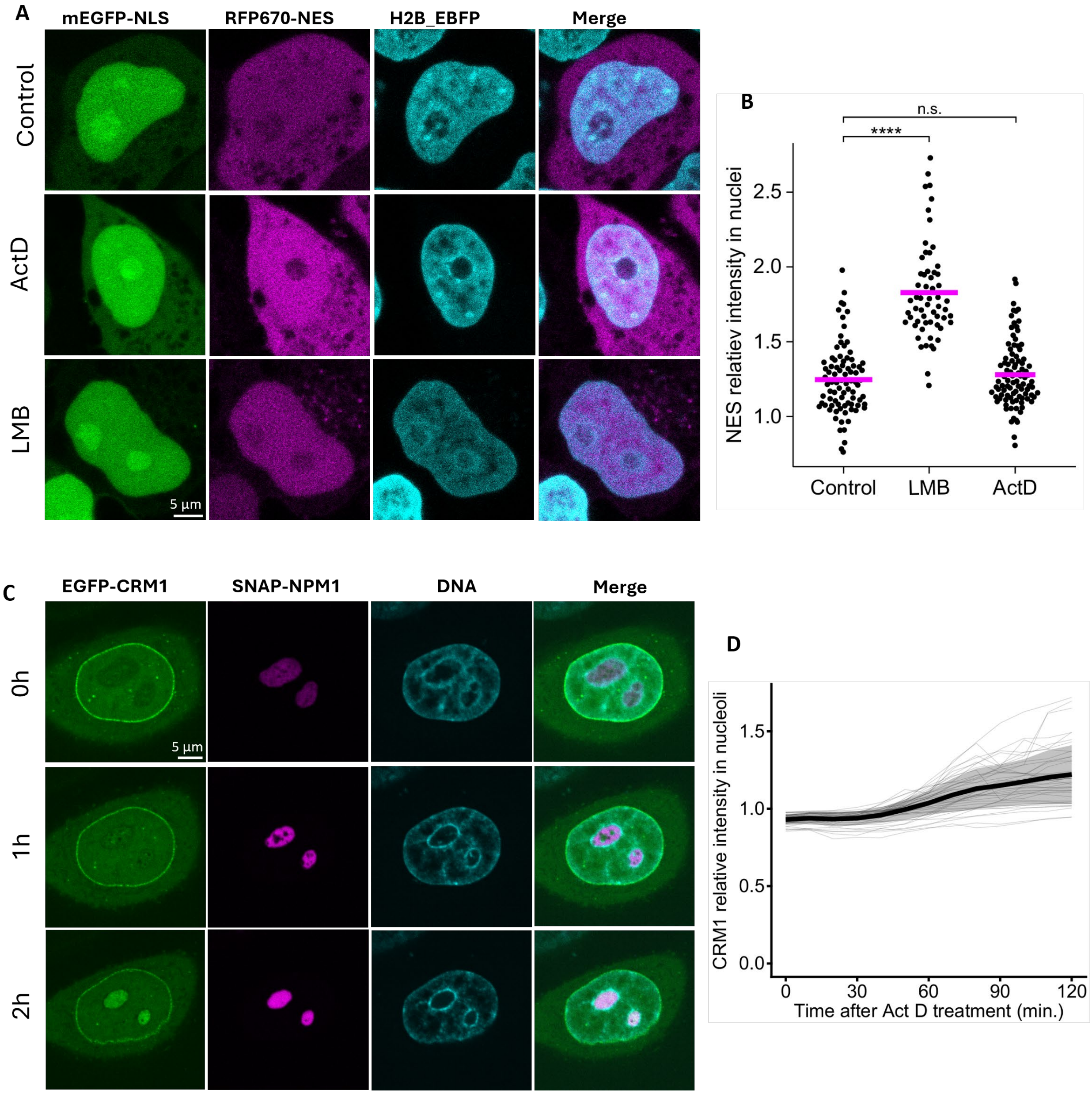


**Fig. S10. CRM1 localization and nuclear export under stress induced by ActD.**

(**A**) EGFP-CRM1 was overexpressed in HeLa cells labeled with SNAP-NPM1, which were treated with 5 nM ActD and imaged by Olympus iXplore SPIN SR with UPLSAPO100XS for 2h with 10 min interval. Scale bar = 5 µm. SNAP-NMP1 (SiR dye labelling), DNA (TMR dye labelling), and EGFP-CRM1. n = 44 cells from 3 biological replicates. (**B**) Mean intensity of the CRM1 signal in the nucleolus was measured and normalized over the signal in nucleoplasm. (**C**) mEGFP-NLS_RFP670-NES nuclear export reporter plasmid was overexpressed in HeLa cells labeled with SNAP-NPM1 and H2B-EBFP. Cells were treated with 5 nM ActD or LMB for 3h. Scale bar = 5 µm. (**D**) Mean intensity of NES signal in the nucleus was measured and normalized over the signal in the cytoplasm. Asterisks denote significance levels determined using the Kruskal–Wallis test followed by Dunn's post hoc test (****P < 0.0001; n.s., not significant). n = 84 cells (Control), n = 59 cell (LMB), n = 94 cells (ActD) from 3 biological replicates.

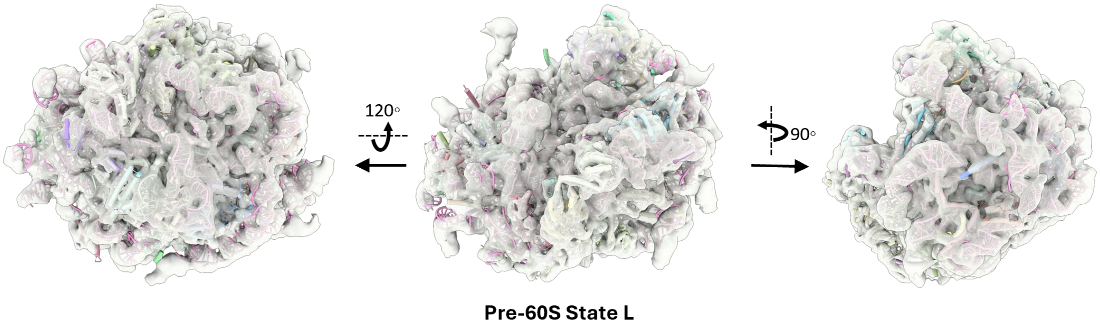


**Fig. S11. Comparison of pre-60S state L and human pre-60S state K3.**

Molecular model of human pre-60S state K3 (PDB 8flc) fitted into the EM map of pre-60S state L.

**Table S1. Human pre-ribosomes datasets detailed across conditions**

| Condition | Untreated | | ActD | | Total | |
|---|---|---|---|---|---|---|
| Magnification | 26000x | | 26000x | | | |
| Voltage (kV) | 300 | | 300 | | | |
| Electron exposure (e-1/Å2) | ~140 | | ~140 | | | |
| Defocus range (µm) | -3 to -7 | | -3 to -5 | | | |
| Pixel size (Å/pixel) | 3.425 | | 3.425 | | | |
| Cell number | 58 | | 28 | | 86 | |
| Tomogram number | 212 | | 93 | | 305 | |
| Total particles | 264799 | | 124584 | | 389383 | |
| Complex | # | % | # | % | Å | EMDB |
| SSU processome state A’ | 20645 | 7.8 | 96 | 0.8 | 8.8 | |
| SSU processome state A | 21782 | 8.2 | 125 | 1.0 | 8.5 | |
| SSU processome state preA1 | 11995 | 4.5 | 350 | 2.8 | 9.3 | |
| SSU processome state preA1$_{exo}$ | 4054 | 1.5 | 21 | 0.2 | 11.4 | |
| SSU processome state postA1$_{exo}$ | 1919 | 0.7 | 15 | 0.1 | 14.1 | |
| SSU processome state postA1 | 6835 | 2.6 | 231 | 1.9 | 10.8 | |
| Pre-60S state A | 8040 | 3.0 | NA | NA | 10.9 | |
| Pre-60S state B | 801 | 0.3 | NA | NA | 19.4 | |
| Pre-60S state C | 2453 | 9.0 | 898 | 7.2 | 12.0 | |
| Pre-60S state D | 32805 | 12.4 | 13511 | 10.8 | 7.7 | |
| Pre-60S state E | 9963 | 3.8 | 3706 | 3.0 | 8.9 | |
| Pre-60S state F | 13143 | 5.0 | 1415 | 1.1 | 8.4 | |
| Pre-60S state G* | 4618 | 1.7 | 1935 | 1.6 | 9.5 | |
| Pre-60S state G | 33942 | 12.8 | 8018 | 6.4 | 7.6 | |
| Pre-60S state H | 45670 | 17.2 | 19661 | 15.8 | 7.1 | |
| Pre-60S state I$^{pre}$ | 7265 | 2.7 | NA | NA | 8.8 | |
| Pre-60S state I$^{post}$ | 11086 | 4.2 | NA | NA | 8.8 | |
| Pre-60S state J | 21378 | 8.1 | 14379 | 11.5 | 7.1 | |
| Pre-60S state K | 3509 | 1.3 | 24587 | 19.7 | 7.3 | |
| Pre-60S state K$_{CRM1}$ | 2896 | 1.1 | 17807 | 14.3 | 7.1 | |
| Pre-60S state L | NA | NA | 17829 | 14.3 | 7.8 | |
